# Earthworm Invasion reduces above-belowground Biodiversity and Ecosystem Multifunctionality

**DOI:** 10.1101/2025.11.05.686452

**Authors:** Olga Ferlian, Nico Eisenhauer, Michael Bonkowski, Marcel Ciobanu, Kenneth Dumack, Lee E. Frelich, Edward A. Johnson, Bernhard Klarner, Benjamin Rosenbaum, Jörg-Alfred Salamon, Madhav P. Thakur, Lise Thouvenot, Tesfaye Wubet, Malte Jochum

## Abstract

Global change alters abiotic and biotic conditions across the globe with unprecedented consequences for the functional integrity of affected ecosystems. Many belowground ecosystems are subject to an underestimated aspect of global change: the invasion of earthworms. While we know that earthworm invasion can impact the physical, chemical, and biological properties of invaded ecosystems, it remains poorly understood how these changes are interconnected and how they concurrently affect the overall functioning of an ecosystem. Unfortunately, most studies addressing global-change impacts focus on a very limited number of environmental variables, taxa, and ecosystem functions. Here, we collected data on six environmental variables, ten functional groups of microbes, plants, and animals, and 16 ecosystem functions from four forests in the USA and Canada, well-known hotspots of earthworm invasion. We used a multi-step Structural-Equation Modeling approach to disentangle the direct and indirect effects of earthworm invasion on environmental conditions and the ecosystem multidiversity and multifunctionality of invaded forests at different levels of resolution. Our analysis revealed that earthworm invasion reduced total multidiversity (combined microbial, plant, and animal multidiversity). Ecosystem multifunctionality was reduced via a combination of direct and indirect effects, with the latter involving both effects, mediated by altered environmental conditions and by total multidiversity. When resolving total multidiversity into taxon-specific multidiversity indices for microbes, plants, and animals, the above-mentioned effects of earthworm invasion on multifunctionality via multidiversity disappeared. Invasion effects on single ecosystem functions differed in their mediators but were net negative across the board. Given these differences across the differently-resolved analyses, the results suggest that a whole-ecosystem perspective is paramount to comprehensively understanding the impacts of biological invasions. Combining a multi-step analytical design with multiple biodiversity indices and multiple ecosystem functions assessed simultaneously, our study represents the most complete assessment of the mechanisms and ecosystem-level consequences of earthworm invasion to date.

## Introduction

Anthropogenic activities accelerate global change, thereby threatening Earth’s ecosystems (Díaz et al. 2018, Balvanera et al. 2019, Isbell et al. 2023). The resulting loss of biodiversity often leads to a decline in ecosystem functions (Hooper et al. 2005, Isbell et al. 2015, Duffy et al. 2017), which, in turn, are the basis for the ecosystem services humanity depends on (Cardinale et al. 2012, Isbell et al. 2017). To date, most research on this global change- biodiversity-functioning nexus has either focused on single taxonomic groups and trophic levels, on single ecosystem functions, or both (but see Soliveres et al. 2016, Eisenhauer et al. 2023). Moreover, most studies on the ecosystem consequences of biodiversity change use random biodiversity-loss scenarios (Bruelheide et al. 2014). However, global change typically leads to non-random changes in biodiversity and community structure (Duffy et al. 2017, Giling et al. 2019), which are likely to have significant ecosystem-scale consequences (Soliveres et al. 2016, Beaumelle et al. 2020). Therefore, if the main biodiversity drivers for a given location or ecosystem type are known, we should consider non-random biodiversity- change and its inherent taxonomic and trophic complexities when studying community- and ecosystem-level processes (Seibold et al. 2018), to gain a holistic understanding of how global change affects ecosystem multifunctionality (Balvanera et al. 2006, Stevens and Tello 2014, Maxwell et al. 2016, Giling et al. 2019).

Ecosystem multifunctionality (hereafter: multifunctionality), the ability of an ecosystem to simultaneously support multiple ecosystem functions, has become a key indicator of functional change in biodiversity-ecosystem functioning research (Maestre et al. 2012, Byrnes et al. 2014, Schuldt et al. 2018). Due to inherent trade-offs in the expression of functional traits among species, multifunctionality is typically expected to increase with species diversity. Since the individual functional contributions differ between species and functional groups, and show different degrees of overlap, we still lack a comprehensive understanding of the key drivers of the overall functioning of an ecosystem. Past studies have found that ecosystem multifunctionality requires a greater number of species, including rare ones, than is typically the case for individual ecosystem functions (Hector and Bagchi 2007, Isbell et al. 2011, Byrnes et al. 2014). In comparison to multifunctionality, combining biodiversity information across multiple taxonomic groups and trophic levels into a synthetic index of multidiversity, is far less common (Allan et al. 2014, Delgado-Baquerizo et al. 2020). However, integrating across both multidiversity and multifunctionality is essential for comprehensively assessing the link between biodiversity change and ecosystem integrity, since individual functions are typically connected to particular trophic and taxonomic groups (Allan et al. 2015, Soliveres et al. 2016, Eisenhauer et al. 2019b). Research suggests that when impacts on multidiversity and multifunctionality are more pronounced than effects on individual taxonomic groups or ecosystem functions, conservation strategies should adopt more comprehensive approaches that encompass a broader range of species and functional processes (Guerra et al. 2022, Li et al. 2024). Despite the powerful potential provided by joining these approaches to comprehensively assess the consequences of global change on entire communities and ecosystem functioning, few studies have simultaneously considered both multidiversity and multifunctionality indices (but see e.g. Soliveres et al. 2016). This is likely due to the obvious challenges underlying the comprehensive assessment of multiple taxa across trophic levels and multiple ecosystem functions at a reasonable scale.

While the responses of aboveground ecosystem functions to global change and biodiversity loss have been comprehensively studied, belowground biodiversity-ecosystem functioning research has received less scientific attention (but see Heemsbergen et al. 2004, Wagg et al. 2014, Delgado-Baquerizo et al. 2020). This is particularly true for studies linking multi-taxon biodiversity to ecosystem functions (Guerra et al. 2020). Due to their rich biodiversity harbouring ∼59% of all species on Earth and their capacity for carbon storage, soil ecosystems play a critical role in mitigating global change and sustaining a functioning biosphere (Bardgett and van der Putten 2014, Crowther et al. 2019, FAO 2020, Anthony et al. 2023). Their biodiversity contributes to a multitude of essential ecosystem functions and services related to, for example, nutrient cycling, plant growth, and water filtration (Bardgett and van der Putten 2014, FAO 2020). Unfortunately, global change threatens soil biodiversity and, with it, above and belowground multifunctionality, for example via surface sealing, deforestation, pollution, or extreme weather events (Orgiazzi et al. 2016, FAO 2020, Phillips et al. 2024). Given their high biodiversity, their importance for maintaining ecosystem multifunctionality, and their vulnerability under global change, soils represent a crucial, yet understudied research focus in global-change ecology and key stressors for these important ecosystems might have been largely overlooked.

Biological invasions in soil ecosystems remain an underestimated dimension of global change (Phillips et al. 2024), despite evidence that belowground invaders like earthworms can fundamentally alter biodiversity and ecosystem functioning (Murphy and Romanuk 2014, Balvanera et al. 2019, Jochum et al. 2022). As ecosystem engineers, earthworms restructure soils through burrowing, mixing, and litter decomposition (Jones et al. 1994, Frelich et al. 2019), and can become highly successful invaders in systems where they are functionally novel to native communities (Wardle et al. 2011). In northern North America, the postglacial reintroduction of European earthworms has triggered widespread shifts in soil properties (Ferlian et al. 2020). These changes include altered soil pH, moisture, and nutrient availability, alongside cascading direct and indirect effects on plants, soil microbes, and soil invertebrates (Craven et al. 2017, Ferlian et al. 2018, 2020, Jochum et al. 2021). Plants are primarily affected through modified seedbed composition and resource availability (Nuzzo et al. 2015, Craven et al. 2017), while microbial communities respond to altered substrate inputs and accelerated decomposition pathways (Ferlian et al. 2018). Soil invertebrates are most strongly affected because the removal and homogenization of the leaf-litter layer threatens their primary habitat, refuge, and food resources, while simultaneously disrupting the microclimatic conditions of the soil-litter interface on which they depend (Brown 1995, Ferlian et al. 2018). At the same time, a substantial proportion of soil invertebrates feed on soil microbes, which may further amplify the effects of earthworm-induced changes (Thakur and Eisenhauer 2015). Although recent (meta-)studies document these impacts, it remains unclear how earthworm-driven shifts in biodiversity and environmental conditions influence ecosystem multifunctionality, given their concurrent effects across interconnected functions. Focusing on single functions may therefore underestimate cumulative impacts, trade-offs, and emergent changes in overall ecosystem functioning.

While observational field studies on earthworm invasion make it difficult to identify causal relationships and specific drivers for functional and biodiversity change (Ferlian et al. 2018), structural equation modeling (SEM) can help disentangle direct invasion effects from those mediated by biodiversity and environmental changes by simultaneously testing multiple interconnected hypotheses (Eisenhauer et al. 2015, Delgado-Baquerizo et al. 2020). Here, we tested the direct and indirect effects of earthworm invasion on ecosystem multidiversity and multifunctionality in four northern North American forests using an SEM approach. We combined six environmental variables with the biodiversity of ten functional groups of microbes, plants, and animals, and 16 measured ecosystem functions. We employed a multi- level SEM approach where we first assessed direct and indirect effects of earthworm invasion on soil environmental properties, multidiversity, and multifunctionality (referred to as aggregated SEM hereafter). In the second step, we zoomed in further and extended these SEMs by resolving the biodiversity of microbes, plants, and animals into separate multidiversity indices (each combining the taxonomic richness of multiple functional groups; referred to as taxon-resolved SEM hereafter). In the final step, we individually explored, for the 16 single ecosystem functions, to what extent their responses to invasive earthworms were mediated by either changes in biodiversity or environmental properties. Overall, we hypothesised that earthworm invasion disrupts resident organism communities and, consequently, decreases multidiversity, likewise directly and indirectly through altered environmental properties (H1). We further hypothesised that earthworm invasion decreases multifunctionality, and this effect is at least partly mediated via reduced multidiversity and altered soil environmental properties (H2). Third, we expected the negative effects of earthworm invasion on biodiversity to be strongest for soil animal multidiversity compared to soil microbial and plant multidiversity (H3), as soil animals experience both direct and indirect effects with the latter cascading up through the soil food web. Finally, we hypothesized earthworm invasion to have predominantly negative effects on single ecosystem functions, with variation in the underlying pathways, such that microbe-related functions are primarily driven by microbial multidiversity, while other functions are governed by their respective biotic or abiotic drivers (H4).

## Material and Methods

### Study site

Data were collected in four northern North American forests located in Alberta, Canada, and in Minnesota, USA. The three Canadian forests (northern bank of Barrier Lake, Alberta: 51.035 N, 115.065 E; 1450 m a.s.l., hereafter called Barrier North; southern bank of Barrier Lake, Alberta: 51.015 N, 115.073 E; 1380 m a.s.l., hereafter called Barrier South; and Bull Creek Hills, Alberta: 50.400 N, 114.554 E; 1300 m a.s.l., hereafter called Bull Creek) are situated in the Canadian Rocky Mountains, Kananaskis Valley, southwest Alberta (Barrier Lake Field Station, University of Calgary). The Canadian forests are dominated by trembling aspen (*Populus tremuloides*) and balsam poplar (*Populus balsamifera*), with a dense understorey and an Orthic Grey Luvisol soil. They are characterized by short, dry summers and cold winters with the soil being frozen between November and March, a mean annual precipitation of 625 mm, and a soil temperature of 3.8°C (Eisenhauer et al. 2019). The US forest is located near St. John’s Abbey in Collegeville, central Minnesota (45.573 N, 94.398 W; 350 m a.s.l.; hereafter called St. John’s). It is dominated by sugar maple (*Acer saccharum*), American basswood (*Tilia americana*), and Northern Red and White oak (*Quercus rubra*, *Q. alba*) on loamy sand Udipsamments (or Orthic Regosols) with a humid continental, cold temperate climate. The annual precipitation is 776 mm, and the mean annual temperature is 7°C (Eisenhauer et al. 2019a).

In the studied regions, only single native earthworm species survived the last glaciation period and are now present in low densities (Hendrix and Bohlen 2002, Csuzdi et al. 2017). However, previous assessments of earthworms in glacial refugia in our study area reported no native taxa (Hilton and Reynolds 1983). The assessed regions have a long history of earthworm-invasion research with clearly-documented invasion fronts (e.g. Hale et al. 2005, Eisenhauer et al. 2007, Straube et al. 2009), which was one of the reasons for their selection. To verify the invasion status and direction before the set-up of sampling plots in the four forests, we assessed earthworm distributions along a transect following the assumed direction of invasion by digging with a spade and by using a modified mustard-extraction method (Gunn 1992, Eisenhauer et al. 2007, Straube et al. 2009). In addition, the detailed historical information available for the sites from three decades of published studies helped to minimise mistaking background site-level variation for true invasion effects.

### Study design

In Barrier North and South and in St John’s forests, we set up plots between July and September 2016, whereas in Bull Creek forest, we set up plots in August 2017. Where possible, we set up highly-invaded plots in areas where all three ecological earthworm groups, epigeic, endogeic, and anecic, were present (Bouché 1977) and low-invaded plots where no or only low densities of epigeic earthworms were present at the same time taking care of the proximity of the two areas and, thus, similarity of abioic and biotic conditions. Any two plots had a minimum distance of 20 m. In total, we set up twenty plots of 1 x 1 m in each forest, with ten plots in the low-invaded and ten plots in the highly-invaded area of each forest. Each plot was subdivided into four quadrants of 0.5 x 0.5 m for different types of samples. In addition, adjacent to each plot, we defined a further area of 0.3 x 1 m where undisturbed soil cores for DNA sequencing were taken.

### Assessment of environmental properties and soil functions

In the first quadrant, we took three 5-cm-diameter soil cores to a depth of 10 cm, mixed them to a composite sample, and stored them in a plastic bag in a cooling box for transport. Composite samples were sieved through a 2-mm sieve and homogenised. Subsamples were then used for measuring several environmental properties.

For measurements of soil pH, 12 g of fresh soil from the composite sample were dried at room temperature and, subsequently, solved in 25 ml 0.01 M CaCl_2_ solution, shaken, left for at least one hour, and measured with a pH-meter (Orion Star A211, Thermo Scientific, Dreieich, Germany). For details on measurements of soil carbon and nitrogen content, gravimetric soil water content, soil aggregate stability, and soil bulk density, see Appendix S1: Section S1. Litter mass was assessed by hand-collecting all litter from the second quadrant, drying it at 60°C for 72 h and weighing it (in dry mass in g per quarter m^2^).

### Assessment of soil microbial communities

For measurements of soil microbial activity (as basal respiration – BAS, and microbial biomass carbon – Cmic), subsamples of 5 g fresh weight were taken from the composite sample and analysed using an O_2_-microcompensation apparatus (Scheu 1992) and following the protocol in Ferlian et al. (2017). The biomasses of soil bacteria, arbuscular mycorrhizal fungi, saprotrophic/ectomycorrhizal fungi, and of plant material in soil were quantified using phospholipid fatty acid (PLFA) analysis. Fatty acids were extracted from the composite soil sample as described in Frostegård et al. (1993). See Appendix S1: Section S1 for details. For DNA-sequencing of soil bacteria, fungi, and protists, three 10 cm deep soil cores (2 cm in diameter) were taken from the adjacent 0.3 x 1 m subplot. For details, see Appendix S1: Section S1. Bioinformatics were conducted as described in Ferlian et al. (2024). From amplicon sequence variants (ASV) and operational taxonomic units (OTU), we calculated species richness. Calculations were based on rarefied bacterial (44000 reads per sample), fungal (5324 reads per sample) and protist (5700 reads per sample) datasets per sample.

### Assessment of plant communities

Vegetation cover of each 1 x 1 m plot was assessed prior to all sampling activities, estimating the proportion of projected area on the soil surface by each plant species separately from the top-down perspective in steps of 5%. For plant biomass assessment, plants were harvested from the second quadrant by cutting all plant shoots at the soil surface with scissors and transferring them into paper bags. The plant material was sorted to species level per plot, assigned to functional groups (herbs, grasses, legumes, woody plants), dried at 60°C for 72 h, and weighed. Based on these data, we calculated plot-specific functional group biomass as well as plant-species-richness values.

Plot-specific canopy openness was assessed by taking pictures with a cell phone (iPhone 6+) and a clipped-on fisheye lens facing to the sky perpendicularly to the soil surface on a tripod at a height of 1.4 m. Pictures were processed with the WinSCANOPY software (Regent Instruments Inc., Quebec, QC, Canada) to calculate canopy openness (in %).

### Assessment of soil faunal communities

Soil micro- (nematodes), meso- and macrofauna were sampled, extracted, identified, and diversity indices and biomasses were calculated. Details can be found in the Appendix of Jochum et al. (2021). Data was scaled to an area of 1 m^2^ (at a depth between 0 and 10 cm). Due to missing data, microfauna (nematode) richness and biomass were not available for one and two plots (different forests), respectively.

### Assessment of earthworm communities

To verify the assignment of our plots to the low- and highly-invaded areas within each forest, and to gain a continuous predictor variable for earthworm-invasion intensity, we assessed earthworm communities. For details on the method of earthworm sampling, the twelve earthworm species found, and earthworm community abundance, biomass, richness, and group richness, see Appendix in Jochum et al. (2021), respectively.

### PCA of environmental properties

We conducted a Principal Component Analysis (PCA) to identify the main dimensions of the six environmental properties measured at the plot level (R package ‘FactoMineR’ from Lê et al. 2008) in order to fit them into the SEM and calculated contributions of variables to PC axes (Appendix S1: Figure S1). Environmental properties were defined as variables that are not ecosystem functions (process-based definition, e.g. in Odum 1969) and are typically directly influenced by earthworm activity. The scores of the resulting principal components 1 (PC1, 57.6%, representing mainly variables canopy openness [24.0%], bulk density [23.8%], vegetation cover [21.7%], pH [17.0%], and litter mass [6.7%]; see Fig S1) and 2 (PC2, 15%, representing soil water content [67.1%] and litter mass [16.1%]) were used as “Environmental 1” and “Environmental 2” variables in the SEMs respectively (see below).

### Calculation of multidiversity indices

To represent the biodiversity of the overall communities as well as that of higher-order taxonomic groups, namely microbes, plant, and animals, we calculated multidiversity as the proportional species richness at the total community level as well as at the group level (Allan et al. 2014, Delgado-Baquerizo et al. 2020). Specifically, microbial multidiversity included bacteria, fungi, and protist richness; plant multidiversity included grass, herb, legume, and woody species richness; and animal multidiversity included macrofauna, mesofauna, and microfauna taxonomic richness. Total community multidiversity included all ten groups.

### Calculation of multifunctionality from ecosystem-function clusters

We used 16 measurable variables representing ecological functions or key ecosystem processes, including microbial, plant, and animal biomass, soil C and N pools, microbial respiration, and soil aggregate stability. Following previous studies (Maestre et al. 2012, Lefcheck et al. 2015), we jointly considered strict ecosystem functions and key processes or properties as ecosystem functions. Variables were grouped into six clusters to account for interdependencies (Manning et al. 2018): microbial, plant, animal, aggregate stability, soil C, and soil N clusters (Appendix S1: Table S2). For plot-level averaged multifunctionality, each variable was standardized to 0–1 and weighted according to its cluster size (e.g., 0.2 per function in a five-variable cluster). We then calculated the weighted mean using the R package multifunc (Byrnes et al. 2014), omitting its internal standardization. Because each of the six clusters had a total weight of 1, values were divided by six to obtain clustered multifunctionality on a 0–1 scale. Two plots lacked microfauna data, resulting in a sample size of 77 for multifunctionality analyses. We also calculated threshold-based multifunctionality (Byrnes et al. 2014). For each function, the mean of the six highest values defined its maximum, and thresholds of 20% and 70% were applied. For each plot, we summed the cluster-specific weights of functions exceeding each threshold. Thus, values could be non-integer (e.g., 1.2). Unless otherwise stated, “multifunctionality” refers to averaged, clustered multifunctionality. See Appendix S1: Section S1 for a more detailed description.

### Assessing pairwise relationships between earthworm invasion, multidiversity indices, and multifunctionality

As a first statistical step, we tested the total effect of earthworm invasion, represented by earthworm biomass, on total, microbial, plant, and animal multidiversity indices. Earthworm biomass was used as a proxy for invasion impact to avoid introducing intercorrelated variables into the SEMs (see Appendix in Jochum et al. 2021). Because earthworm species vary substantially in body size, biomass provides a more ecologically meaningful measure of functional impact than abundance and has been successfully used in previous studies (Craven et al. 2017, Jochum et al. 2021). Earthworm biomass (g m⁻Z + 1) was log10-transformed to meet model assumptions. We fitted four separate linear mixed-effects models, using earthworm biomass as the fixed-effect predictor and each multidiversity index as the response. We then tested the direct effects of each multidiversity index on averaged multifunctionality in four separate models, followed by a model testing the total effect of earthworm biomass on multifunctionality. Forest identity (n = 4) was included as a random factor in all models because forests differed in sampling time and were not the primary focus. Forest identity explained considerable variation, indicating that effects differed among forests. However, the forest showing the strongest or weakest effect varied among response variables, suggesting that no single forest disproportionately drove the results (Appendix S1: Figure S2). We therefore focused on general patterns integrating diversity indices, ecosystem functions, and study sites. See Appendix S1: Section S1 for a more detailed description.

### Structural equation modelling

To test the direct and indirect effects of earthworm invasion on environmental properties, multidiversity, ecosystem functions, and multifunctionality, we used a SEM approach in piecewiseSEM (version 2.3.0.1, Lefcheck 2016). Specifically, we used a three-step approach starting with both multidiversity and multifunctionality in their most aggregated form, i.e., total multidiversity and multifunctionality (aggregated SEM, n=77, see Appendix S1: Figure S3 for initial SEM structure and Table S1 for hypotheses behind arrows). In this step, we allowed earthworm invasion (earthworm biomass) to directly impact variables “Environmental 1” and “Environmental 2” (Principal components 1 and 2 from the PCA, respectively, see Appendix S1: Figure S1), total multidiversity, and multifunctionality (Appendix S1: Figure S3). The environmental variables were allowed to directly affect total multidiversity and multifunctionality and total multidiversity could also directly impact multifunctionality. This step was intended to study the overall relationships among earthworm invasion, environmental, diversity, and function data. However, given the highly- aggregated nature of the diversity and function data used here, we subsequently complemented this SEM with higher-resolution data. In the second step, we replaced total multidiversity by more-highly resolved multidiversity indices, i.e., the multidiversity indices of microbes, plants, and animals, respectively, providing each of these variables with the same SEM links as total multidiversity in the previous step (taxon-resolved SEM, n=58, see Appendix S1: Figure S4). This step was intended to provide a better understanding of which functional group relates to earthworm impacts up to multifunctionality. In the third step, we kept the three multidiversity indices separate but replaced multifunctionality by the 16 single functions in 16 separate SEMs (fully-resolved SEM, n=60, except for mesofauna biomass, where n=58, see Appendix S1: Figure S4). This step was intended to provide an in-depth understanding of how earthworm invasion impacts the single processes in the four forests and which functional groups are primarily involved in controlling each function. In all SEMs, the two environmental variables shared a correlated error, and in all taxon-resolved and fully- resolved SEMs, there were correlated errors between each pair of the three multidiversity indices involved.

As our initial SEMs were fully saturated (all paths set, Appendix S1: Figure S3, S4), piecewiseSEM could not estimate coefficients of SEM model fit. To be able to estimate model fit, we therefore removed the weakest (lowest std. estimate) non-significant predictor of our target variable, multifunctionality (in fully-resolved SEMs, the target function), from our fully-saturated models to obtain SEMs with model fit parameters Fisher’s C and the related p-value. These values, together with standardized path coefficients, as well as marginal and conditional RZ values are presented in our SEM results figures.

For the aggregated SEM step, we included all forests. Since microbial richness data was entirely lacking for one forest and we thus had to remove it from the taxon-resolved and fully-resolved SEM steps (as microbial multidiversity is a standalone variable in there), we tested if removing that forest from the aggregated SEM resulted in any major differences to our main results. As this SEM did not majorly differ from the one including all forests (Appendix S1: Figure S5 and Table S4), we concluded that the three steps of our SEM approach are comparable. We further explored the sensitivity of our results by testing alternative aggregated SEMs using the 20% and 70% multifunctionality thresholds (Appendix S1: Figure S6 and Table S4). We used R version 4.4.0 “Puppy Cup” for the PCA, mixed-effects models, and SEM analyses.

## Results

### Relationships between earthworm invasion, multidiversity, and multifunctionality

Based on our linear mixed-effects models, earthworm invasion (log_10_) significantly decreased total (Fig. 1a; Appendix S1: Table S3; p < 0.001, R^2^ = 0.21, R^2^ =0.61), plant (Fig. 1c, p = 0.001, R^2^ = 0.04, R^2^ = 0.74), and animal multidiversity (Fig. 1d, p < 0.001, R^2^ = 0.47, R^2^ = 0.56), thereby partly supporting the hypothesis that multidiversity decreases with earthworm invasion (H1). Total multidiversity (Fig. 1e, p < 0.001, R^2^ = 0.32, R^2^ = 0.54), plant (Fig. 1g, p < 0.001, R^2^ = 0.25, R^2^ = 0.54), and animal multidiversity (Fig. 1h, p < 0.001, R^2^ = 0.15, R^2^ = 0.66), in turn, were significantly positively correlated to ecosystem multifunctionality. Overall, earthworm invasion significantly decreased ecosystem multifunctionality (Fig. 1i, p < 0.001, R^2^ = 0.25, R^2^ = 0.73) partly supporting our hypothesis H2. As indicated by the marginal and conditional R^2^-values, forest identity contributed considerably to the explained variance, especially to that of plant multidiversity as affected by earthworm invasion. These pairwise relationships can, however, only provide a first step towards understanding the complex interplay between earthworm invasion, multidiversity, and ecosystem multifunctionality. Therefore, more in-depth analyses were conducted next.

**Fig. 1.**
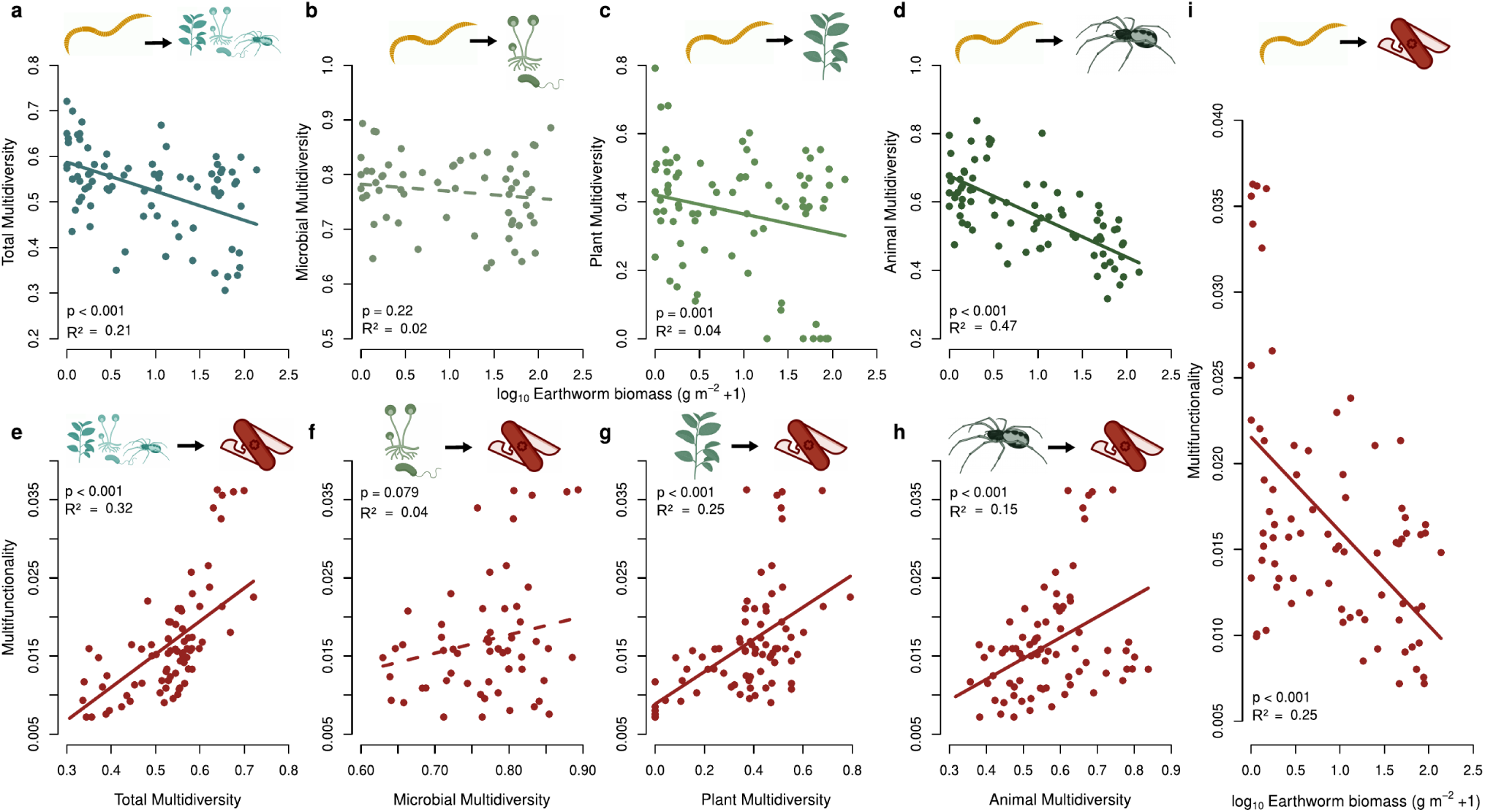
| Pairwise relationships between earthworm invasion, multidiversity indices, and multifunctionality. Effects of earthworm invasion (as represented by earthworm biomass, upper row, log_10_-transformed) on total (a), microbial (b), plant (c), and animal multidiversity (d) and (lower row), of total (e), microbial (f), plant (g) and animal (h) multidiversity on multifunctionality, and (i, log_10_-transformed) earthworm invasion on multifunctionality. Panels show solid and dashed regression lines for significant and non-significant effects, respectively, based on linear mixed-effects models with forest identity as a random factor. Multidiversity and multifunctionality responses are shown in shades of green and red, respectively. Each dot represents a plot in one of the four observational forest sites (upper row: n=80, except for microbes, where n=60, lower row and panel i: n=77, except for microbes, where n=58). See Appendix S1: Table S3 for further model coefficients. Icons created in https://BioRender.com.

### Direct and indirect earthworm-invasion effects

Our aggregated SEM (Fig. 2) revealed that the detrimental effect of earthworm invasion on ecosystem multifunctionality was partly mediated by a decrease in total multidiversity and partly via altered environmental properties, namely soil water content and litter mass. As expected, earthworm invasion directly reduced total multidiversity (hypothesis H1). We did not find any evidence for environmental properties mediating the effects of earthworm invasion on total multidiversity. Furthermore, as expected under our hypothesis H2, earthworms reduced multifunctionality. This effect decomposes into a direct negative component (-0.27) and two indirect negative components. The first indirect component acted via soil water content (-0.29) and its effect on multifunctionality (+0.46) combining into a negative indirect effect (-0.13). In addition, another indirect negative effect of earthworms on multifunctionality resulted from a negative impact of earthworms on multidiversity (-0.37) and its knock-on effect on multifunctionality (+0.23) combining into a weaker indirect negative effect (-0.09). In contrast to earthworms, environmental properties other than soil water content (Environmental 1) affected multidiversity (+0.39), and thus also multifunctionality, but this variable was not driven by earthworm invasion. We did not find any cascading effects from earthworm invasion via both environmental properties and multidiversity to multifunctionality. Although our aggregated SEM illustrates two important indirect effects of earthworm invasion on ecosystem multifunctionality, there was a substantial direct effect indicating additional mechanisms neither explained by multidiversity, nor by the environmental variables but instead directly controlled by earthworms. This suggests that these multifunctionality responses might be driven by ecosystem functions directly controlled by earthworms without necessarily involving the environmental properties, such as for example soil-nutrient concentrations, or diversity indices.

**Fig. 2.**
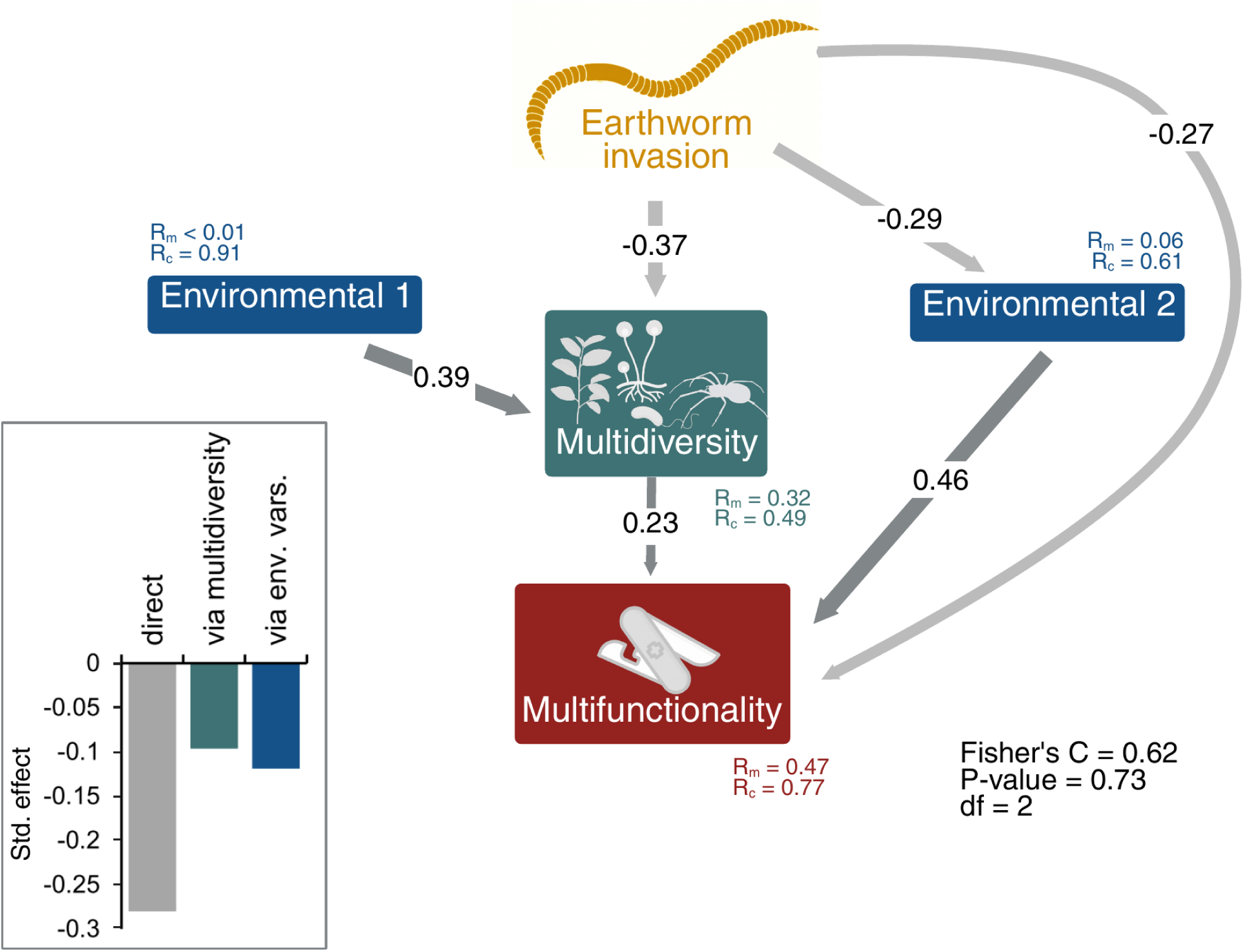
| Structural Equation Model showing the effects of earthworm invasion on aggregated multidiversity and multifunctionality. (n=77). The environmental variables 1 and 2 depict axes 1 and 2 of a Principal Component Analysis (PCA); see Appendix S1: Figure S1 for the PCA biplot). Arrow widths and associated values represent the strength of the relationships (only significant ones, P < 0.05) using standardised path coefficients. Dark and light grey arrows represent positive and negative path coefficients, respectively. The embedded bar plot shows the summarized effects of earthworm invasion on multifunctionality that are either direct, indirect mediated by multidiversity, or indirect mediated by environmental properties. Marginal and conditional RZ values are reported below boxes. Model fit parameters were extracted from piecewiseSEM. Created in https://BioRender.com.

Alternative SEM analyses using a 20% multifunctionality threshold yielded results similar to the averaged multifunctionality approach (Appendix S1: Table S4 and Figure S6). However, a 70% threshold analysis showed no direct effect of earthworm invasion on multifunctionality and no significant relationship between multidiversity and multifunctionality, thereby also eliminating the indirect effects of earthworm invasion on multifunctionality mediated through biodiversity changes.

As shown by the taxon-resolved SEM that separates microbial, plant, and animal multidiversity pathways, earthworm invasion reduced multifunctionality through two pathways (Fig. 3), directly (-0.27) and indirectly via water-related environmental variables (- 0.17). We found that earthworm invasion significantly reduced both plant (-0.18) and more strongly animal multidiversity (-0.64), which additionally had positively correlated residuals (+0.26), but not microbial multidiversity. However, none of these effects had a knock-on effect on multifunctionality, which is in contrast to the fully-resolved SEM. Further, neither of the three multidiversity indices showed indirect responses to earthworm invasion that were mediated by environmental variables. Strong direct effects of earthworm invasion on ecosystem multifunctionality were consistently observed.

**Fig. 3.**
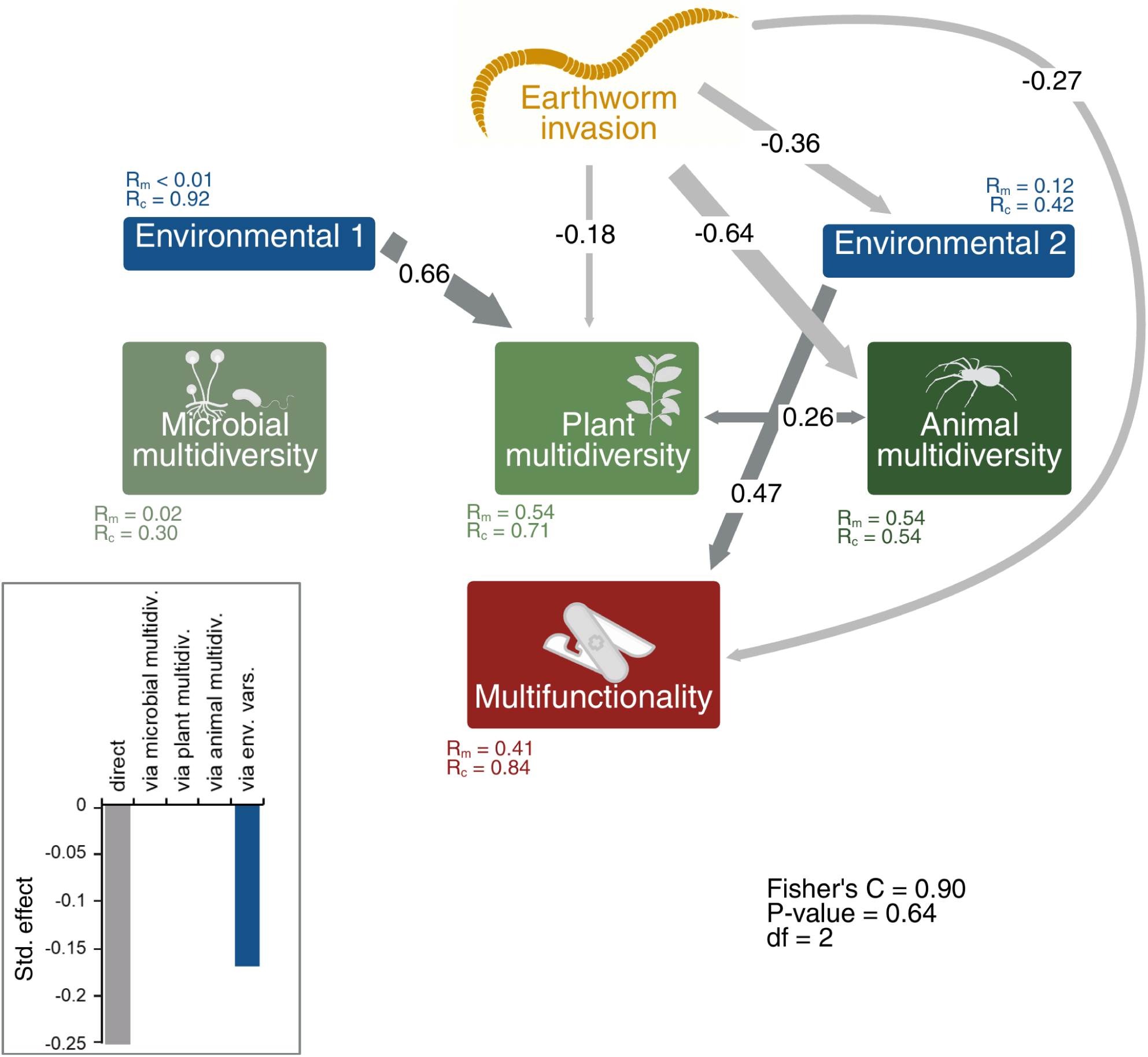
| Structural Equation Model showing the effects of earthworm invasion on taxon- resolved, multidiversity indices as well as multifunctionality. (n=58). The earthworm invasion variable is based on earthworm-biomass data. The environmental variables 1 and 2 depict the axes 1 and 2 of a Principal Component Analysis using six environmental properties (pH, water content, soil bulk density, litter mass, vegetation cover, and canopy openness).

The width of the arrows and associated values represent the strength of the relationships (only significant ones, P < 0.05) using standardised path coefficients. Dark grey arrows represent positive and light grey arrows negative path coefficients. The embedded bar plot shows the summarized effects of earthworm invasion on multifunctionality that are either direct, indirect mediated by microbial, plant, or animal multidiversity, or indirect mediated by environmental properties. Created in https://BioRender.com.

Our detailed analysis of individual ecosystem functions revealed that earthworm invasion significantly affected only half of the functions studied (Fig. 4, Appendix S1: Table S5). Specifically, eight functions showed negative impacts of earthworm invasion (both direct and indirect), while the remaining eight functions remained unaffected. Of those functions affected by earthworm invasion, there was not a single one showing only a direct earthworm effect. Where indirect effects occurred, they were mediated exclusively by either biodiversity changes or environmental alterations, but never by both pathways simultaneously. Among the microbial functions, microbial basal respiration and biomass carbon were decreased by earthworm invasion, whereas the functions related to specific microbial taxa (AMF, fungi, and bacteria) remained unaffected. Earthworm invasion influenced microbial functions through two parallel pathways of approximately equal strength: directly through the earthworms’ presence, and indirectly by earthworms altering environmental properties that subsequently affected microbial processes. Among the plant functions, only woody species biomass was significantly decreased by earthworm invasion, whereas other plant functional groups were not affected. Effects on woody species biomass were mainly mediated via environmental properties and only to a small extent by plant multidiversity. All three animal- related functions were negatively impacted by indirect effects of earthworm invasion.

**Fig. 4.**
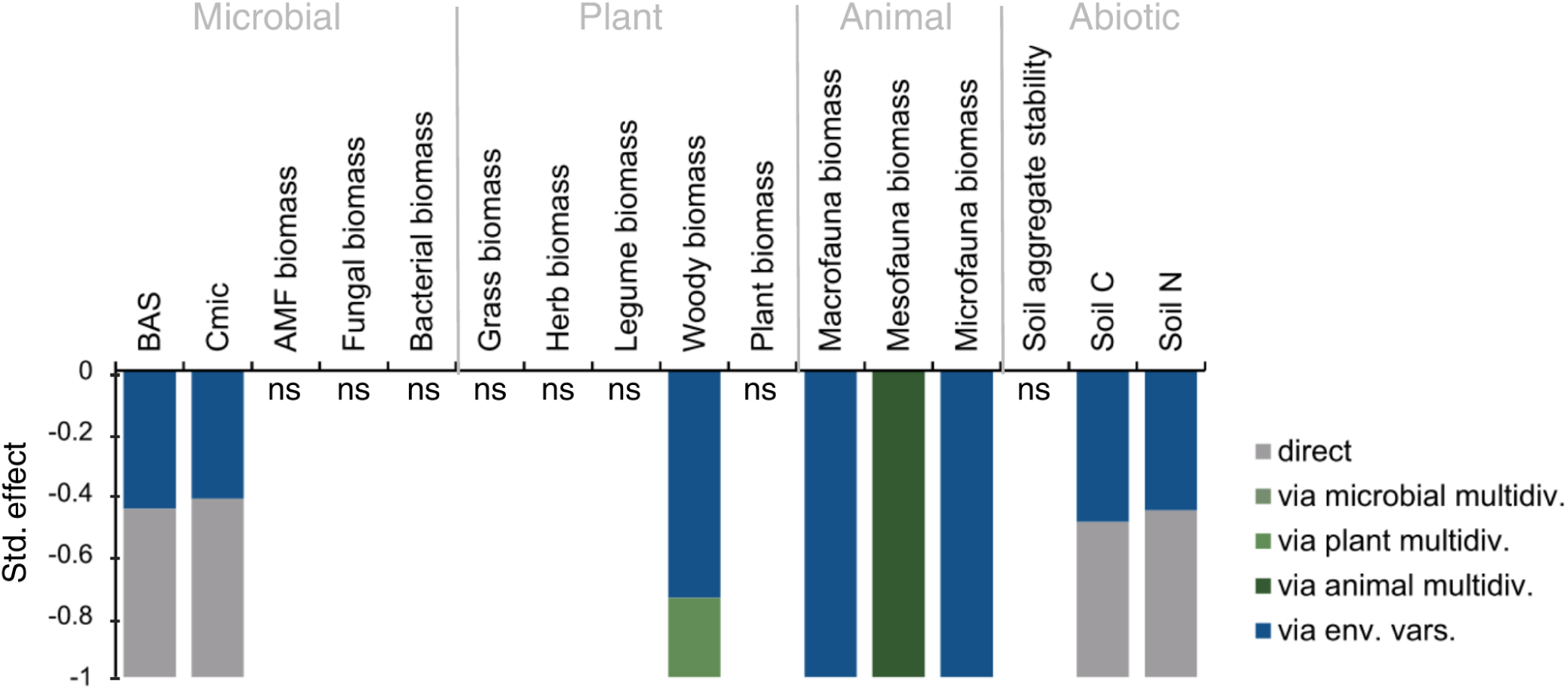
| Barplot of the summarised effects of earthworm invasion on 16 single ecosystem functions. (from 16 separate fully-resolved Structural Equation Models) with the effects being direct, indirect mediated by microbial, plant, and/or animal multidiversity, and/or indirect mediated only by environmental properties. Each bar shows the relative proportion of each of the five paths to the total effect. See Appendix S1: Table S5 for all p-values and standardised path coefficients. BAS: soil microbial respiration, Cmic: soil microbial biomass C, AMF: arbuscular mycorrhizal fungi, C: carbon, N: nitrogen, ns: path not significant. n=60 for all SEMs summarized here, except for the microfauna biomass SEM, where n=58. Created in https://BioRender.com

Notably, macro- and microfauna biomass were exclusively influenced via changes in environmental conditions, whereas mesofauna biomass was affected by shifts in animal multidiversity. Among the three functions in the abiotic cluster, soil carbon and nitrogen, but not soil-aggregate stability, were negatively affected by earthworm invasion, likewise in a direct way as well as mediated through environmental properties. Microbial multidiversity did not mediate any of the indirect earthworm effects in our study. In summary, these results showed varying effects of earthworm invasion on different ecosystem functions partly supporting our hypothesis H4. However, we did not find any evidence for positive effects of the invasion on the 16 assessed functions.

## Discussion

Our study showed that earthworm invasion decreased ecosystem multifunctionality mediated by both reduced multidiversity and altered environmental properties. Resolving total multidiversity into taxon-specific multidiversity indices, the above-mentioned effects of earthworm invasion on multifunctionality via multidiversity disappeared. Further resolving multifunctionality into single ecosystem functions showed that the strength of earthworm effects varied with target ecosystem function. Generally, animal- and plant-related ecosystem functions were most and least affected, respectively.

### Effects of earthworm invasion on multidiversity

Earthworm invasion directly reduced total multidiversity, without mediation by the environmental factors assessed, partially supporting H1. This was unexpected, as earthworms typically influence diversity through their alterations of the soil environment (Eisenhauer et al. 2007). The most obvious impact of earthworm invasion is leaf-litter removal (Hendrix 2006, Suárez et al. 2006), which plays a dual role: for many soil organisms it is a primary food resource as well as their microhabitat (Bohlen et al. 2004, Eisenhauer et al. 2007). Litter mass, however, included via variable Environmental 2 in the PCA, contributed surprisingly little to explain earthworm effects on multidiversity. The weak mediating effect of litter mass may reflect a temporal mismatch, as winters in these forests are relatively long and, at the time of sampling, recently deposited leaf litter may not yet have been substantially processed by earthworms, resulting in only minor differences between invaded and uninvaded sites that consequently did not mediate effects of earthworm biomass on multidiversity indices or multifunctionality. Soil aeration, represented by both, soil bulk density and soil water content, is typically another key soil property commonly affected by earthworm invasion (Ferlian et al. 2020, Edwards and Arancon 2022), and, although it was a significant driver of multifunctionality in our study, soil aeration showed little impact on multidiversity. The lack of indirect earthworm-invasion effects on multidiversity may stem from opposing influences of the measured environmental properties on different taxa, which could have been masked in our aggregated SEM analysis (Ferlian et al. 2018). An alternative explanation for the absence of indirect earthworm effects on multidiversity via environmental variables is that focusing on a single earthworm variable (here: invasive-earthworm biomass) as a proxy for invasion may not capture the full complexity of earthworm impacts. Our aggregated SEM showed that environmental variables (Environmental 1: primarily canopy openness, bulk density, pH, vegetation cover, and litter mass) influenced total multidiversity, but earthworm invasion had no significant effect on these environmental variables. This suggests that other aspects of the earthworm community - such as species composition, richness, abundance, the relative importance of ecological groups (epigeic, endogeic, anecic), or body-size distribution - may further regulate environmental conditions which, in turn, could indirectly affect multidiversity (Craven et al. 2017). While we focused on earthworm biomass as a common proxy for earthworm invasion (Eisenhauer 2010, Craven et al. 2017, Jochum et al. 2022), future research should further disentangle the relative contribution of such invasive earthworm-community properties on the biodiversity and ecosystem performance of invaded systems.

### Effects of earthworm invasion on ecosystem multifunctionality

Earthworm invasion decreased multifunctionality which is in line with previous findings exploring the effects of invasive earthworms on individual ecosystem functions (Bohlen et al. 2004, Huang et al. 2010, Ferlian et al. 2020). As hypothesised (H2), these effects were mediated in part by the negative effects on multidiversity and altered environmental properties. Both these drivers were similarly important in our study suggesting two distinct mechanisms how earthworms can shift habitat conditions (‘ecosystem engineer’, Jones et al. 1994, Wu et al. 2025) and multitrophic biodiversity, thus threatening the overall integrity of invaded ecosystems. In general, it has to be considered that we used a set of six environmental properties that entered the PCA and that mainly soil water content represented the second axis that drove the environmentally-mediated effects in the SEMs. This was unexpected, since drivers like mechanical disturbance and litter mass often dominate earthworm effects (Migge 2001, Bohlen et al. 2004; here, only litter mass was involved in significant effects via a minor contribution to Environmental 2). Soil water content had a greater impact than the other properties on the measured ecosystem functions, such as biomasses of microbes and soil fauna, that are typically soil moisture-dependent (Yeates 1979, Drenovsky et al. 2004, Kardol et al. 2011). Different mechanisms have been reported by which earthworm activity can change soil-water regimes, for example, by increasing water-infiltration rates in most soil types (Pérès et al. 1998, Capowiez et al. 2014) or decreasing rates in others (Frelich et al. 2006). Further, soil evapotranspiration has been found to increase due to litter removal and exposition of bare soil, which decreases soil water content (Bohlen et al. 2004). The other environmental properties also affected multidiversity and thus multifunctionality, but we did not find support of earthworm biomass driving these environmental properties. This does, however, not necessarily mean that they do not mediate earthworm-invasion effects on multifunctionality – but might simply indicate that one of the most powerful and simple proxies for earthworm invasion, earthworm biomass, might not represent these additional effects of earthworm invasion (Craven et al. 2017, Jochum et al. 2022).

Besides the indirect effects of earthworm invasion on multifunctionality, we found a considerable direct effect, which combines variability that could not be explained by our measured mediating variables with potential direct earthworm impacts – represented by those functions that can be directly influenced by earthworms. For example, other metrics related to the studied biota than the assessed richness, such as functional diversity, specific traits or species composition and identity, may be affected by earthworm invasion to a similar extent and result in earthworm effects on multifunctionality that cannot be explained by richness making them direct earthworm effects in our SEMs. Moreover, functions like elemental content may be directly related to earthworm invasion without being mediated by multidiversity nor environmental properties (Postma-Blaauw et al. 2006), which may even bypass litter mass effects that contributed to the PCA axis Environmental 2.

The alternative SEM version with threshold-based multifunctionality indices can also further contribute to our understanding of earthworm-invasion impacts on multifunctionality. Using multiple thresholds captures how multifunctionality responds across a range of performance levels, revealing trade-offs, synergies, and the robustness of ecosystem functions more accurately than any single threshold could (Byrnes et al. 2014, Liu et al. 2019). We found that using a 20% threshold led to significant results and results highly comparable to those of our aggregated SEM analysis. However, with a threshold level of 70%, the number of paths between earthworm-invasion effects and multifunctionality decreased, and the link between multidiversity and multifunctionality disappeared. This suggests that earthworm invasion influences the ability of ecosystems to maintain low-to-moderate levels of multiple functions, but not high levels of multiple functions simultaneously. In other words, earthworms may shift baseline functioning across several processes, yet they do not enhance (or consistently suppress) the system’s capacity to sustain functions at high performance levels. This pattern points to weak but widespread effects across functions rather than strong, coordinated effects on top-end ecosystem performance. Liu et al. (2019) showed the opposite pattern, namely that earthworm effects increased with multifunctionality thresholds, but this study did not assess earthworm effects in an invasion context. More studies assessing (invasive) earthworm impacts on multifunctionality, ideally combining different standard multifunctionality approaches, will help to further contextualize these results in the future.

### Microbial, plant, and animal multidiversity as mediators of earthworm-invasion effects

Both our mixed-effects models and the taxon-resolved SEM showed that earthworm invasion did not significantly affect microbial multidiversity. For our microbial diversity data derived from DNA-sequencing, we sampled the top 0-10 cm soil layer which included the litter as well as the mineral horizon. A former meta-analysis showed that effects of earthworm invasion on microbial communities are negative in the organic and positive in the mineral layer (Ferlian et al. 2018). Thus, combining topsoil layers in the present study could have averaged out potential opposing effects across the soil profile. In comparison, effects of earthworm invasion on animal multidiversity were strong which confirms our third hypothesis and suggests that effects on faunal communities seem to be rather consistent across soil horizons and/or that soil animals are more prone to earthworm activities (Ferlian et al. 2018). This is attributable to the fact that they rely on the physical and macroaggregate structure of the soil more than microbes that occupy a much smaller microhabitat. Moreover, decomposer taxa, such as Collembola and oribatid mites may compete with earthworms for food resources to a certain extent (Eisenhauer 2010), thus explaining why they might respond more strongly to earthworm invasion. Interestingly, we did not find any indirect effects of earthworm invasion on the different multidiversity indices that were mediated by environmental properties, which partly rejects our third hypothesis. This finding may be due to an unexplained proportion of the effects mediated by environmental properties that were not covered by the two PC axes, such as that related to mineralisation or leaching (Hale et al. 2005, Resner et al. 2015, Ferlian et al. 2020). Interestingly, also none of the taxon-specific multidiversity indices mediated earthworm-invasion effects on multifunctionality. This might suggest that a multidiversity index integrating across different taxonomic groups is necessary to capture the overall effects of earthworm invasion on ecosystem functioning. Weak effects on individual taxa may not be detectable within single indices, but can accumulate to significant impacts when integrated into a composite (total) multidiversity metric (Duffy 2002, Wang et al. 2019).

### Resolving multifunctionality into single functions

Resolving both total multidiversity into taxon-specific multidiversity indices and multifunctionality into single ecosystem functions revealed that half of the functions were significantly decreased by earthworm invasion, whereas the other half remained unaffected. This only partly supports H4, expecting to find largely negative effects. Functions related to soil animals, to general microbial biomass, and to soil carbon and nitrogen contents decreased with earthworm invasion. Similar to the strong earthworm effects on animal multidiversity, animal-related ecosystem functions may be more strongly affected by earthworm invasion due to multiple direct and indirect effects cascading towards higher trophic levels compared to the other organisms. However, it was unexpected that the microbial functions related to single microbial groups were not significantly affected. But the results also showed that there are still direct effects pointing to the fact that variables that were not included in the analyses may have played a role. Among the plant-related functions, only woody species biomass (which included only shrubs and tree seedlings/saplings in our study) significantly decreased with earthworm invasion, while the biomass of other plant functional groups like grasses and herbs remained unaffected. This result was unexpected, since former studies mainly showed strong positive effects on grasses (Drouin et al. 2016, Alexander et al. 2022, Thouvenot et al. 2024a) as well as negative effects on herbaceous plants due to their contrasting nutrient- uptake strategies and drought tolerance (Craven et al. 2017, Alexander et al. 2022). However, the decrease in woody plants observed is consistent with the results from Nuzzo et al. (2009) and Thouvenot et al. (2024a), who both found a decrease in the cover of woody plant species with earthworm invasion. The significant negative effect in woody species may be due to the burrowing effects on seed germination and/or due to altered soil nutrient concentrations which rather benefit resource-acquisitive growth strategies in plant functional groups like grasses or herbs (Eisenhauer and Scheu 2008, Craven et al. 2017). Disentangling multifunctionality into single functions further showed that effects on some functions, like woody biomass and the animal-related ones, were entirely mediated by our measured environmental properties as well as multidiversity indices. Our analysis revealed that the patterns clearly visible in the aggregated SEM linking earthworms to multifunctionality via environmental properties and total multidiversity were not detected in the more resolved SEMs. Invasive earthworms might not significantly affect every single taxonomic group or each of the 16 ecosystem functions in our analysis but they do affect their multidiversity and the ability of invaded ecosystems to simultaneously maintain multiple ecosystem functions. Surprisingly, and in contrast to our fourth hypothesis, the single functions were not driven by their corresponding biotic and abiotic pathways. Although it has been shown that animal- diversity and -biomass effects are often linked (Schneider et al. 2016), our results indicate that this link is rather weak in the context of earthworm invasions and that rather abiotic drivers (or further unknown direct effects in SEM) play an essential role.

## Conclusions

Our study provides strong evidence that earthworm invasion decreases multidiversity and multifunctionality of northern North American forest soils. The effects were mainly mediated by significant reductions in soil water content and the combined diversity of multiple organism groups, including microbes, plants, and animals, suggesting a major threat to native biodiversity. Previous meta-studies focused on certain taxa or trophic groups and single ecosystem functions (Cameron et al. 2016, Craven et al. 2017, Ferlian et al. 2018) potentially underestimating the overall effects of earthworm invasion on forest ecosystems. This study represents the first attempt to comprehensively assess these effects, with the biodiversity of multiple higher-order taxa and multiple ecosystem functions explored at the same time within the same ecosystems. This simultaneous assessment of multiple variables is a key advancement of our approach building on previous meta-analyses that, while able to add many factors, are typically limited by the various spatial and temporal scopes of their included data sets. Our study thus enhances our mechanistic understanding of biological invasions and draws a more complete picture of their causes and consequences to fully understand their complex, cascading impacts on multiple ecosystem components and functions. While earthworms are essential drivers of nutrient cycling (Wu et al. 2025) and key facilitators of global food production in their native ecosystems and in agricultural systems (Fonte et al. 2023), our work stresses their detrimental role as invaders at the ecosystem scale.

## Acknowledgements

We thank Svenja Haenzel, Anja Pasemann, and Farah Lodhawalla for help with ECOWORM coordination, Ulrich Pruschitzki, Tom Künne, Julius Quosh, Ian Macdonald, Anja Zeuner, Simone Cesarz, Michelle Ives, Adrienne Cunnings, Artur Stefanski, Sofia Pipp, Ian Macdonald, and numerous student helpers for support in the field and lab, and the Barrier Lake Field Station (Kananaskis, Alberta, Canada) and St. John’s University (Minnesota, USA) for accommodation and support. We acknowledge funding by the European Research Council under the European Union’s Horizon 2020 research and innovation program (grant no. 677232 to NE) and the European Union HORIZON-MISS-2022-SOIL-01-03 project Integrating Soil Biodiversity to Ecosystem Services (SOB4ES, grant no. 101112831 to MB). Further support came from the German Centre for Integrative Biodiversity Research (iDiv) Halle-Jena-Leipzig and the Gottfried Wilhelm Leibniz Prize, funded by the Deutsche Forschungsgemeinschaft (DFG, German Research Foundation) – FZT 118, 202548816; Ei862/29-1. We used ChatGPT (OpenAI) to refine the language of this manuscript.

## Author contributions

NE developed the general ECOWORM project design and received the funding. OF and MPT led the data collection and lab analyses. OF, MB, MC, KD, LF, BK, JAS, MPT, LT, and TW processed microbial, plant, and animal samples and performed the related biodiversity and function assessments. OF and MJ collated the data, designed and performed all statistical analyses, and drafted the paper. BR helped design the multi-step SEM analysis. All authors contributed to interpreting the results and writing the manuscript.

