## Appendix for "Earthworm Invasion reduces above-belowground Biodiversity and Ecosystem Multifunctionality"

**Appendix S1**

**Earthworm invasion reduces above-belowground biodiversity and ecosystem multifunctionality**

**Journal: Ecology**

Olga Ferlian[1,2], Nico Eisenhauer[1,2], Michael Bonkowski[3], Marcel Ciobanu[4], Kenneth Dumack[3], Lee E. Frelich[5], Edward A. Johnson[6], Bernhard Klarner[7], Benjamin Rosenbaum[1,8], Jörg-Alfred Salamon[9], Madhav P. Thakur[10], Lise Thouvenot[1,2], Tesfaye Wubet[11,12], Malte Jochum[1,2,13]

### Section S1: Materials and Methods

*Soil carbon and nitrogen content*

A subsample of 10 g fresh soil was dried at 60°C for 72 h. Soil was ground with a ball mill, subsequently dried for another 24 h, transferred into tin capsules, and weighed. Analyses were performed using an elemental analyser (Vario EL II, Elementar Analysensysteme GmbH, Hanau, Germany). Carbon and nitrogen contents are given as relative mass proportion of the element (in %) per sample dry mass.

*Soil water content*

Gravimetric soil water content was measured from a subsample of 5 g fresh weight that was weighed before and after drying at 60°C for 72 h to calculate the relative mass proportion of water in fresh soil.

*Soil bulk density*

For calculations of soil bulk density, one further soil sample per treatment and forest was taken using a soil corer of 5 cm diameter and 10 cm depth. Samples were dried at 105°C for 24 h and weighed. Soil bulk density was calculated as dry mass per volume (g $cm^{-3}$).

*Soil-aggregate stability*

For measurements of soil-aggregate stability, one soil sample per plot was gently taken in the first quadrant with a hand shovel without compacting the soil, transferred into a 50 ml Falcon tube, and cooled. In the lab, 4 g of air-dried aggregates were weighed into small sieves and sieved with a wet-sieving apparatus (Eijkelkamp Soil & Water, Giesbeek, the Netherlands) according to Kemper and Rosenau (1986). We calculated the proportion of water-stable aggregates (in %) using soil mass before and after sieving.

*Biomass of soil microbial taxa*

The biomasses of soil bacteria, arbuscular mycorrhizal fungi, saprotrophic/ectomycorrhizal fungi, and of plant material in soil were quantified using phospholipid fatty acid (PLFA) analysis. Fatty acids were extracted from the composite soil sample as described in Frostegård et al. (1993). Lipid fractions were saponified, methylated, and washed as described in the protocols for the Sherlock Microbial Identification System (MIDI Inc., Newark, USA). The resulting fatty acid methyl esters were analysed with a gas chromatograph (Clarus 680, PerkinElmer, Waltham, USA; carrier gas helium; flame ionization detector; SP-2560 capillary column 100 m × 0.25 mm i.d., 0.2 µm film thickness), and subsequently fatty acid biomasses were calculated. In the analysis, we used marker fatty acids indicative of bacteria (i15:0, a15:0, i16:0, i17:0, cy17:0, cy19:0, 16:1ω7), arbuscular mycorrhizal fungi (neutral lipid fraction, 16:1ω5), and saprotrophic/ectomycorrhizal fungi (18:2ω6,9; Ruess and Chamberlain (2010)).

*Richness of soil microbes using DNA-sequencing*

For DNA-sequencing of soil bacteria, fungi, and Protists, three 10 cm deep soil cores (2 cm in diameter) were taken from the adjacent 0.3 x 1 m subplot, sieved (2 mm) directly in the field, transferred to 15 ml Falcon tubes, and immediately stored on dry ice. Soil samples for DNA sequencing were only taken in three of the four forests (not in Bull Creek) due to technical reasons. Before taking each sample, all corers, tools, and gloves were washed and sterilised with 70% ethanol to avoid any contamination with DNA across plots. Samples were frozen at -80°C in the lab until further processing. Genomic DNA was extracted using a PowerSoil DNA Isolation Kit (MO BIO Laboratories Inc., Carlsbad, California, USA) following the manufacturer's protocol, with some modifications (Ferlian et al. 2024). DNA yields were checked and further processed as described in Ferlian et al. (2024). The sequencing data of the fungal and bacterial amplicons were deposited in the National Center for Biotechnology Information (NCBI) Sequence Read Archives (SRA) under the BioProject number PRJNA1001019. Sequencing data of the protists were deposited at the European Nucleotide Archive (ENA) under the accession number ERS2039495 (SAMEA104421553).

*Nematodes*

A subsample of 25 g from the composite sample was used for nematode extraction with the modified Baermann method (here, referred to as soil microfauna, Cesarz et al. 2019). Extracted nematodes were transferred to 4% hot formalin and counted (total abundance of nematodes). From each sample, 100 individuals were randomly selected and identified to genus (adults and most of the juveniles) or family level (juveniles) following Bongers (1988). Plot-level nematode taxonomic richness (hereafter referred to as richness) was calculated from these data. Nematode body masses (genus- and family-specific) were retrieved from the Nemaplex database (http://nemaplex.ucdavis.edu) on 19 August 2019 (for details, see Jochum et al. 2021) and scaled to an area of 1 m2 (at a depth between 0 and 10 cm) using soil bulk density data. Due to missing data, microfauna (nematode) richness and biomass were not available for one and two plots (different forests), respectively.

*Soil mesofauna*

Soil mesofauna was sampled by taking one 5-cm-diameter soil core to a depth of 10 cm per plot in the first quadrant and transported to the lab. Mesofauna was extracted from the cores following a heat-extraction method (Jochum et al. 2021). Extracted animals were transferred to 70% ethanol and identified to species (Collembola and Oribatida) or higher-order taxon level (other Acari) (Christiansen and Bellinger 1998, Weigmann 2006). Plot-level mesofauna richness was calculated from these data. Mesofauna biomass was calculated by measuring up to five individuals per morphospecies-plot combination for body length and afterwards using length-mass regressions to estimate fresh body masses as described in Jochum et al. (2021). Data was scaled to an area of 1 m2 (at a depth between 0 and 10 cm). Due to missing data, microfauna (nematode) richness and biomass were not available for one and two plots (different forests), respectively.

*Soil macrofauna*

Soil surface-active macrofauna was sampled with a combination of sieving and hand-sorting on the third and fourth quadrant within each plot. First, litter and organic soil from one quadrant was collected by hand, sieved into a box through a 2 cm-mesh sieve, and then hand-sorted for animals. Simultaneously, the bare plot area was screened for 10 min, and appearing macrofauna was caught with forceps. Afterwards, the same procedure was applied to the other quadrant. All animals were stored in 70% ethanol and identified to species level or the highest taxonomic level possible and plot-level macrofauna richness was calculated from these data. Mesofauna and earthworms were removed from the macrofauna dataset for the analysis. Macrofauna biomass was calculated in the same way as for mesofauna (see above).

*Earthworms*

We assessed earthworm communities in the second quadrant within each plot immediately after harvesting the vegetation using a combination of digging and hand-sorting (upper 10 cm of the soil) as well as the modified mustard extraction method (Gunn 1992, Jochum et al. 2021) collecting earthworms for 30 min. Earthworms were stored in 70% ethanol and identified to species level. We assessed total earthworm fresh biomass per plot (g $m^{-2}$, hereafter referred to as earthworm invasion) by weighing and summing individual fresh masses.

*Calculation of multifunctionality from ecosystem-function clusters*

We used 16 variables that all are measurable proxies of ecological processes or ecosystem performance, such as microbe, plant, and animal biomasses as proxies for productivity, energy flow, and trophic functioning, soil C and N pools as proxies for nutrient storage and cycling capacity, microbial respiration as proxy for microbial metabolic activity and soil carbon cycling, and soil aggregate stability as proxy for soil structural function, water regulation, and erosion resistance. This definition of ecosystem function is in line with former studies (Maestre et al. 2012, Lefcheck et al. 2015), but we are well aware that our variables constitute two categories, ecosystem functions in a strict sense and key processes or properties, i.e., ecosystem functions in a wider sense (Maestre et al. 2012). Here, we jointly treat variables from both categories as ecosystem functions. The functions were allocated to six pre-defined clusters (Appendix S1: Table S2) to account for their interdependencies (Manning et al. 2018): cluster 1 (microbial cluster) - BAS, Cmic, AMF biomass, fungal biomass, and bacterial biomass (the latter three derived from PLFA measurements, see above), cluster 2 (plant-related cluster) -

grass biomass, herb biomass, legume biomass, woody biomass (all derived from weighing plant material), and total plant biomass in soil (derived from PLFA measurements), cluster 3 (animal-related cluster) - macrofauna biomass, mesofauna biomass, and microfauna biomass, cluster 4 - soil aggregate stability, cluster 5 - soil C, and cluster 6 - soil N (clusters 4 to 6 called abiotic clusters hereafter).

To calculate plot-level averaged multifunctionality, while also taking the function clusters into account, we first standardised each variable between 0 and 1, then weighted each function with its cluster-specific weight (a cluster with 5 variables means weight 0.2 for each function), and finally applied a standard averaging method (R package 'multifunc', function getStdAndMeanFunctions from Byrnes et al. 2014). This method needed to be slightly adjusted by skipping its built-in standardization to allow for the order of first standardizing and then weighting functions which was crucial for our clustering approach. To account for the clustering approach where the weight of each of the six clusters was 1, the resulting multifunctionality values were then divided by 6, to represent the averaged clustered multifunctionality in relation to the maximum-possible value of 6 reached if all functions were at their maximum of 1. Since two plots lacked microfauna biomass, we could not calculate multifunctionality for these plots hence lowering our sample size for analyses including multifunctionality to 77.

As an alternative multifunctionality method commonly used, we calculated several threshold-based multifunctionalities (Byrnes et al. 2014), again adjusted to our clustering approach. For each function, we averaged the six highest values as the function maximum, then calculated selected threshold values (20%, 70%), and subsequently determined for each plot how many functions were above that threshold value. For all functions above the respective thresholds, we summed their cluster-dependent weights per plot. As a result, this clustered threshold multifunctionality variable does not only include integers (counting 1 for each function above the threshold) but could, for example, take a value 1.2, if two functions are above the threshold where one function is from a single-function cluster and the other one from a cluster with five functions. In this paper, if not clearly noted otherwise, the term "multifunctionality" refers to averaged, clustered multifunctionality.

*Assessing pairwise relationships between earthworm invasion, multidiversity indices, and multifunctionality*

As a first statistical analysis step, we tested the total effect of earthworm invasion, represented by earthworm biomass, on total, microbial, plant, and animal multidiversity indices. We used earthworm biomass as a proxy for earthworm impact to avoid introducing multiple intercorrelated variables into the SEMs (see Appendix of Jochum et al. 2021, for details related to earthworm variables). Moreover, because earthworm species exhibit substantial variation in body size, biomass provides a more ecologically meaningful measure of their functional impact than abundance and it has successfully been used in previous studies on earthworm invasion impacts on plants (Craven et al. 2017) and soil animals (Jochum et al. 2021).

Earthworm biomass (in g $m^{-2}$, plus 1 to allow log-transformation for plots without earthworms) was $\log_{10}$-transformed to meet model assumptions. We used four separate linear mixed-effects models with earthworm biomass as the predictor (fixed effect) and each of the four multidiversity indices as the response variable. Next, we tested the direct effect of each of those multidiversity indices on averaged multifunctionality in four separate linear mixed-effects models with a single multidiversity as the predictor (fixed effect) and averaged multifunctionality as the response. Finally, we tested the total effect of earthworm biomass (as above) as a predictor (fixed effect) on averaged multifunctionality. In all linear mixed-effects models, we used forest identity (n=4) as a random factor. This was particularly important because differences among forests were not the primary focus of the study and, for organizational reasons, the forests were sampled at slightly different time points throughout the summer season. Our analyses revealed that forest identity explained a considerable amount of variation, suggesting that the effects we saw differed between the studied forests. However, the identity of forests with stronger or weaker impacts on our models varied across response variables indicating that no single forest represented an odd one out majorly driving the results of our analysis (Appendix S1: Figure S2). This further highlights that a joint analysis across different sites could help identifying more general patterns than any single forest could have provided. Thus, the focus of the current study was to find general patterns amalgamating diversity indices and ecosystem functions as well as study sites.

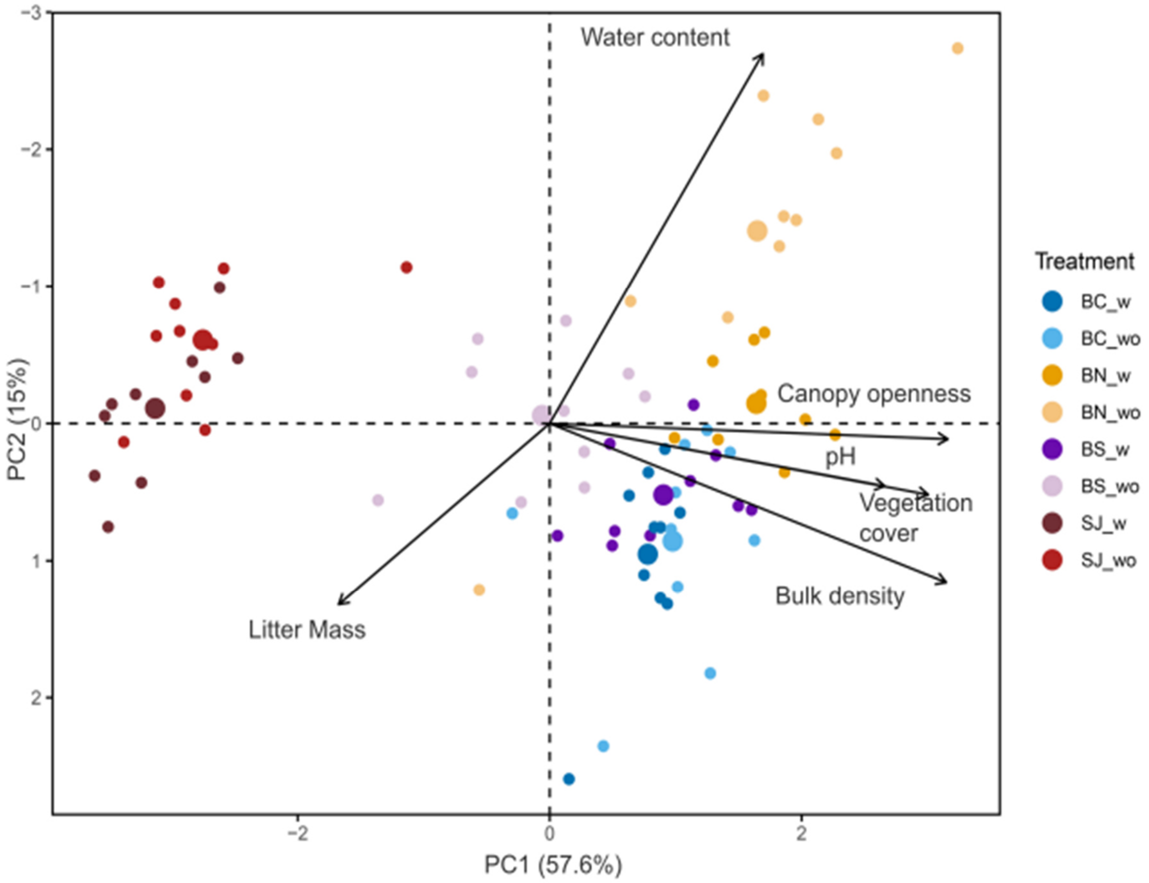


**Fig. S1** Principal Component Analysis (PCA) ordination plot of all environmental variables (represented by the black vectors) and samples (coloured dots) measured in four northern North American forests in 20 plots each. Large dots represent the centroids of each forest and earthworm treatment. w: highly-invaded area, wo: low-invaded area, BC: Bull Creek, BN: Barrier North, BS: Barrier South, SJ: St. John's

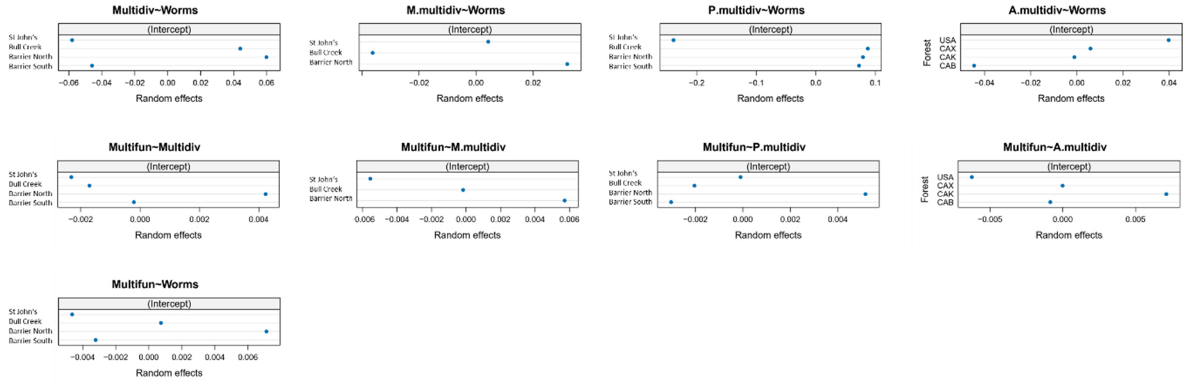


**Fig. S2** Random effects of the pairwise relationships between earthworm invasion, multidiversity indices, and multifunctionality (Fig. 1) for each of the four forests separately(linear mixed-effects models). M.multidiv: microbial multidiversity; P.multidiv: plant multidiversity; A. multidiv: animal multidiversity.

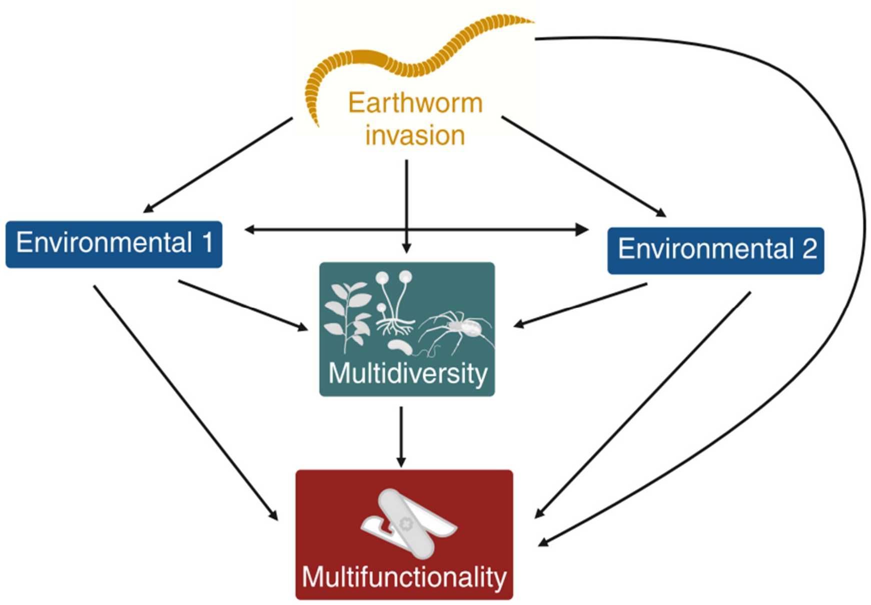


**Fig. S3** The conceptual aggregated structural equation model (SEM) testing the causal relationships among earthworm invasion, environmental properties (represented by the first two principal component axes of six environmental variables), total multidiversity, and multifunctionality. Single-headed arrows represent a hypothesised causal effect among variables, whereas double-headed arrows represent hypothesised correlated errors among variables. Created in https://BioRender.com

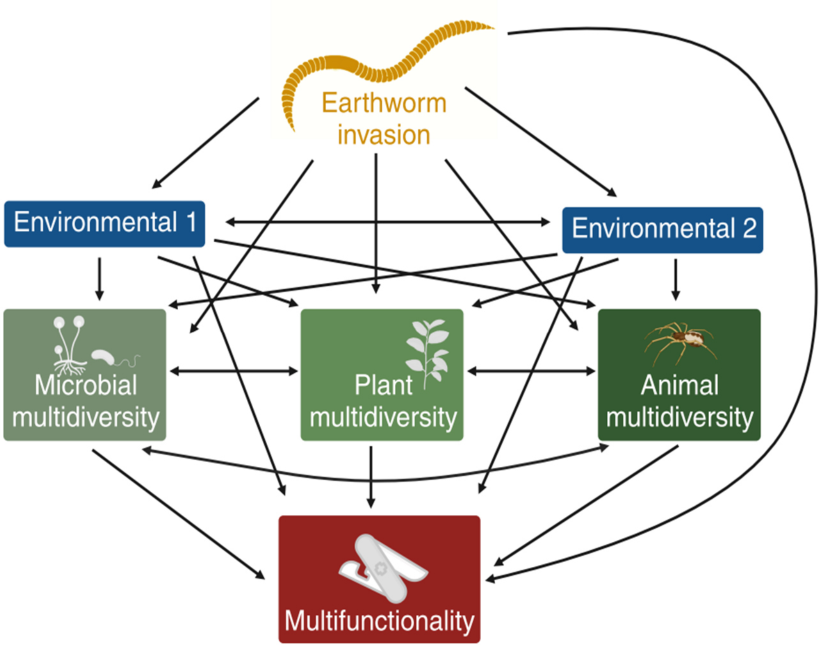


**Fig. S4** The conceptual taxon-resolved structural equation model (SEM) testing the causal relationships among earthworm invasion, environmental properties (represented by the first two principal component axes of six environmental variables), microbial, plant, and animal multidiversity as well as multifunctionality. Single-headed arrows represent a hypothesised causal effect among variables, whereas double-headed arrows represent correlated errors among variables. The same structure was used for the fully-resolved SEMs, simply replacing multifunctionality by the single functions. Created in https://BioRender.com.

Table S1 Predictions for the structural equation model (SEM) paths investigating the relationships between earthworm invasion and (multi)diversity indices and multifunctionality/single ecosystem functions with respective literature support. The three sections of the table refer to the aggregated (Fig. 2), taxon-resolved (Fig. 3), and full-resolved (Fig. 4) SEMs.

| **Predictor** | **Response** | **References** |
|---|---|---|
| Earthworm invasion | Environmental variables | Capowiez et al. 2014, Ferlian et al. 2020 |
| | Multidiversity | See second section (single diversity indices) |
| | Multifunctionality | Constán-Nava et al. 2015 |
| Environmental variables | Multidiversity | Schuldt et al. 2015 |
| | Multifunctionality | Hu et al. 2021, Zhao et al. 2024 |
| Multidiversity | Multifunctionality | Delgado-Baquerizo et al. 2020 |
| Earthworm invasion | Microbial multidiversity | Ferlian et al. 2018, Ferlian et al. 2024 |
| | Plant multidiversity | Craven et al., 2017 |
| | Animal multidiversity | Ferlian et al. 2018, Jochum et al. 2021, 2022 |
| Environmental variables | Microbial multidiversity | Zheng et al. 2019, Hu et al. 2021 |
| | Plant multidiversity | Hu et al. 2021 |
| | Animal multidiversity | Schuldt et al. 2015 |
| Microbial multidiversity | Multifunctionality | Delgado-Baquerizo et al. 2020, Hu et al. 2021 |
| Plant multidiversity | Multifunctionality | Meyer et al. 2018, Hu et al. 2021 |
| Animal multidiversity | Multifunctionality | Delgado-Baquerizo et al. 2020 |
| Earthworm invasion | microbial functions | Groffman et al. 2004, Ferlian et al. 2018 |
| | plant functions | Craven et al. 2017 |
| | animal functions | Jochum et al. 2021, 2022 |
| | abiotic functions | Hale et al. 2005, Ferlian et al. 2020 |
| Environmental variables | microbial functions | Malik et al. 2018, Jiang et al. 2024 |
| | plant functions | Neina 2019, Majasalmi and Rautiainen 2020 |
| | animal functions | de Vries et al. 2012 |
| | abiotic functions | Zheng et al. 2019 |
| Microbial multidiversity | microbial functions | Delgado-Baquerizo et al. 2016, Nannipieri et al. 2017 |
| | plant functions | Delgado-Baquerizo et al. 2016 |
| | animal functions | Barnes et al. 2017 |
| | abiotic functions | Delgado-Baquerizo et al. 2016, Zheng et al. 2019 |
| Plant multidiversity | microbial functions | Lange et al. 2015 |
| | plant functions | Tilman et al. 2001, Isbell et al. 2009 |
| | animal functions | Eisenhauer et al. 2011, Zhang et al. 2022 |
| | abiotic functions | Lange et al. 2015 |
| Animal multidiversity | microbial functions | Delgado-Baquerizo et al. 2020 |
| | plant functions | Delgado-Baquerizo et al. 2020 |
| | animal functions | Barnes et al. 2014 |
| | abiotic functions | Delgado-Baquerizo et al. 2020 |

**Table S2** Ecosystem-function clusters used to calculate averaged multifunctionality. For each function, the table shows the cluster it belongs to, the number of functions in that cluster, and the resulting proportional weight of the individual function.

| Function | Cluster ID | No. included functions | Weight |
|---|---|---|---|
| BAS | 1 | 5 | 0.2 |
| $C_{mic}$ | 1 | 5 | 0.2 |
| AMF biomass | 1 | 5 | 0.2 |
| fungal biomass | 1 | 5 | 0.2 |
| bacterial biomass | 1 | 5 | 0.2 |
| grass biomass | 2 | 5 | 0.2 |
| herb biomass | 2 | 5 | 0.2 |
| legume biomass | 2 | 5 | 0.2 |
| woody biomass | 2 | 5 | 0.2 |
| plant biomass | 2 | 5 | 0.2 |
| macrofauna biomass | 3 | 3 | 0.33 |
| mesofauna biomass | 3 | 3 | 0.33 |
| microfauna biomass | 3 | 3 | 0.33 |
| soil aggregate stability | 4 | 1 | 1 |
| soil C | 5 | 1 | 1 |
| soil N | 6 | 1 | 1 |

**Table S3** Results of linear mixed-effects models presented in Fig. 1. For each model, the table shows the predictor and response variables, sample size (n), whether the predictor was transformed to meet model assumptions, slope estimates, approximated lower and upper 95% confidence intervals (using function 'intervals' from R package 'nlme' (Pinheiro et al. 2014), t- and p-value and marginal (only fixed effects) and conditional (fixed and random effects) pseudo $R^2$ values calculated using function 'rsquared' from R package 'piecewiseSEM' (Lefcheck 2016). Forest identity was used as a random intercept factor for all models. Significant p-values ($p < 0.05$) are given in bold.

| Response | Predictor | n | Pred transf. | Slope | Lower | Upper | t-value | p-value | $R^2$ marginal | $R^2$ conditional |
|---|---|---|---|---|---|---|---|---|---|---|
| **total multidiversity** | **EW biomass** | **80** | **log10** | **-0.06** | **-0.08** | **-0.04** | **-6.07** | **<0.001** | **0.21** | **0.61** |
| m. multidiversity | EW biomass | 60 | log10 | -0.01 | -0.03 | 0.01 | -1.24 | 0.22 | 0.02 | 0.29 |
| **p. multidiversity** | **EW biomass** | **80** | **log10** | **-0.06** | **-0.09** | **-0.02** | **-3.37** | **0.001** | **0.04** | **0.74** |
| **a. multidiversity** | **EW biomass** | **80** | **log10** | **-0.12** | **-0.15** | **-0.09** | **-8.72** | **<0.001** | **0.47** | **0.56** |
| **multifunctionality** | **total multidiversity** | **77** | **none** | **0.04** | **0.03** | **0.06** | **6.04** | **<0.001** | **0.32** | **0.54** |
| multifunctionality | m. multidiversity | 58 | none | 0.02 | 0 | 0.05 | 1.79 | 0.079 | 0.04 | 0.52 |
| **multifunctionality** | **p. multidiversity** | **77** | **none** | **0.02** | **0.01** | **0.03** | **3.98** | **<0.001** | **0.25** | **0.54** |
| **multifunctionality** | **a. multidiversity** | **77** | **none** | **0.03** | **0.02** | **0.04** | **5.59** | **<0.001** | **0.15** | **0.66** |
| **multifunctionality** | **EW biomass** | **77** | **log10** | **-0.01** | **-0.01** | **0** | **-7.94** | **<0.001** | **0.25** | **0.73** |

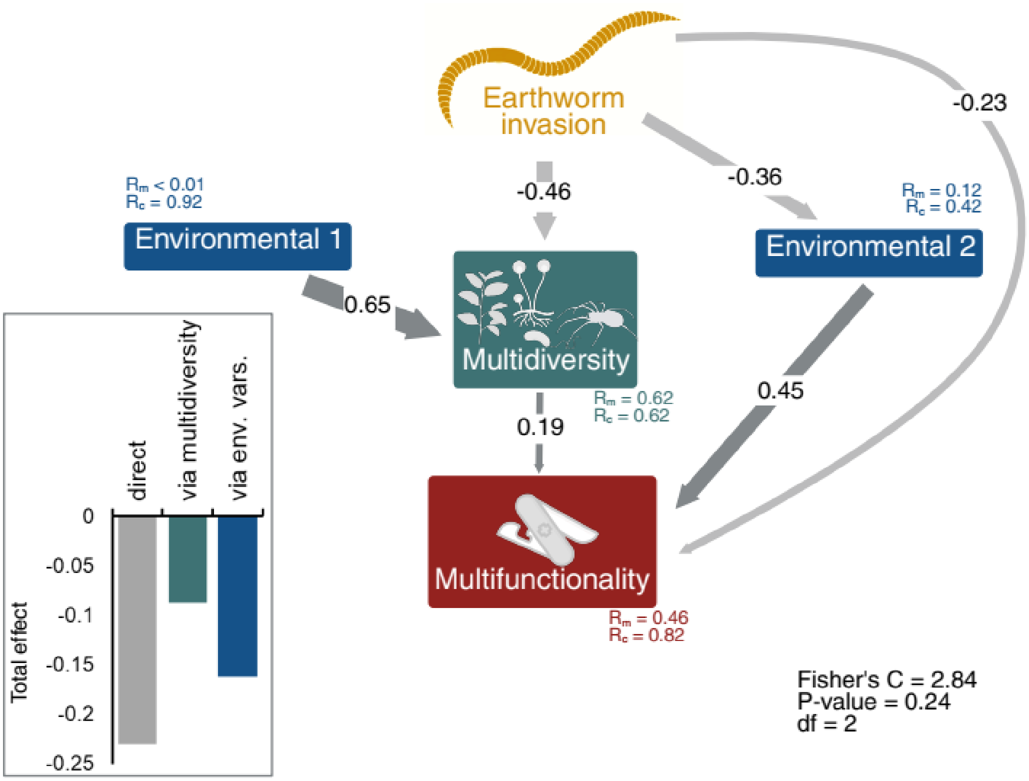


**Fig. S5** Alternative Structural Equation Model showing the aggregated effects of earthworm invasion on multidiversity and multifunctionality with three instead of four forests included (Bull Creek excluded, n=60). The earthworm variable is based on earthworm biomass data. The environmental variables 1 and 2 depict axes 1 and 2 of a Principal Component Analysis (PCA); see Fig. S1 for the PCA biplot). Multidiversity was calculated using averaged proportional species richness per taxon in ten microbial, plant, and animal subgroups. Multifunctionality was calculated from 16 ecosystem functions. Arrow widths and associated values represent the strength of the relationships (only significant ones, $P < 0.05$) using standardised path coefficients. Dark and light gray arrows represent positive and negative path coefficients, respectively. The embedded barplot shows the summarised effects of earthworm invasion on multifunctionality that are either direct, indirect mediated by multidiversity, or indirect mediated by environmental properties. Marginal and conditional $R^2$ values are reported below boxes. Model fit parameters were extracted from piecewiseSEM. Created in https://BioRender.com.

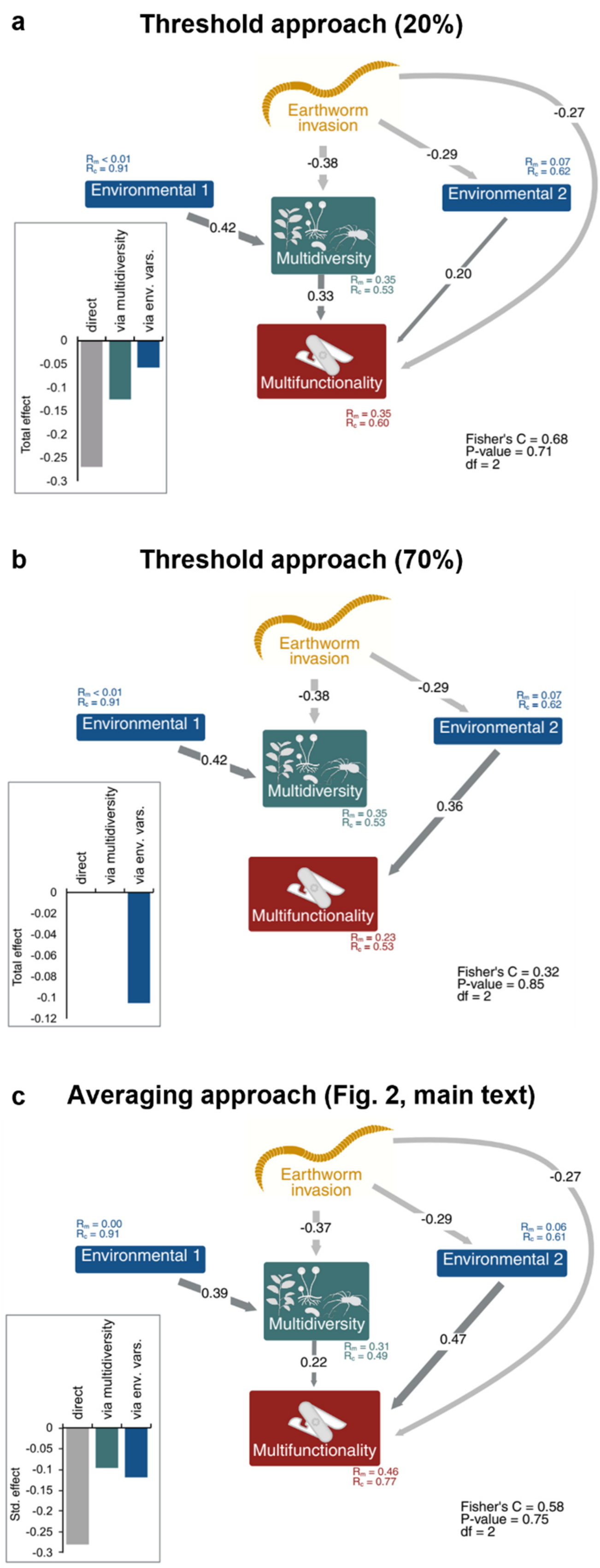


**Fig. S6** Structural Equation Models showing the aggregated effects of earthworm invasion on multidiversity and multifunctionality (n=77) with different approaches for calculating ecosystem multifunctionality: threshold approach 20% (a), threshold approach 70% (b), and averaging approach (c, same as Fig.2, main text). The earthworm variable is based on earthworm biomass data. The environmental variables 1 and 2 depict axes 1 and 2 of a Principal Component Analysis (PCA); see Fig. S1 for the PCA biplot). Multidiversity was calculated using averaged proportional species richness per taxon in ten microbial, plant, and animal subgroups. Multifunctionality was calculated from 16 ecosystem functions. Arrow widths and associated values represent the strength of the relationships (only significant ones, $P < 0.05$) using standardised path coefficients. Dark and light gray arrows represent positive and negative path coefficients, respectively. The embedded barplot shows the summarized effects of earthworm invasion on multifunctionality that are either direct, indirect mediated by multidiversity, or indirect mediated by environmental properties. Marginal and conditional $R^2$ values are reported below boxes. Model fit parameters were extracted from piecewiseSEM. Created in https://BioRender.com.

**Table S4** Results (p-values and standardized path coefficients) of the paths in the Structural Equation Model used in Fig. 2 and in the three alternative models using only three forests (without Bull Creek (BC), Fig. S4), the 20%-threshold approach (Fig. S5a), and the 70% threshold approach (Fig. S5b) for calculating multifunctionality. Env: Environmental variable (PC axis), EW: Earthworm.

| | | Aggregated SEM (see Fig. 2) | | Aggregated SEM without BC | | Aggregated SEM (20% threshold) | | Aggregated SEM (70% threshold) | |
|---|---|---|---|---|---|---|---|---|---|
| Response | Predictor | p | Path coefficient | p | Path coefficient | p | Path coefficient | p | Path coefficient |
| multifunctionality | EW biomass | **<0.001** | **-0.27** | **<0.01** | **-0.23** | **<0.01** | **-0.27** | 0.08 | -0.17 |
| multifunctionality | multidiversity | **<0.01** | **0.23** | **0.04** | **0.19** | **<0.01** | **0.33** | 0.25 | 0.12 |
| multifunctionality | Env2 | **<0.001** | **0.46** | **<0.001** | **0.45** | **0.05** | **0.20** | **<0.01** | **0.36** |
| multidiversity | EW biomass | **<0.001** | **-0.37** | **<0.001** | **-0.46** | **<0.001** | **-0.38** | **<0.001** | **-0.38** |
| multidiversity | Env1 | **0.03** | **0.39** | **<0.001** | **0.65** | **0.02** | **0.42** | **0.02** | **0.42** |
| multidiversity | Env2 | 0.12 | 0.18 | 0.61 | 0.05 | 0.14 | 0.17 | 0.14 | 0.17 |
| Env1 | EW biomass | 0.68 | 0.02 | 0.67 | 0.02 | 0.79 | 0.01 | 0.79 | 0.01 |
| Env2 | EW biomass | **<0.01** | **-0.29** | **<0.01** | **-0.36** | **<0.001** | **-0.29** | **<0.001** | **-0.29** |

**Table S5** Results (p-values and standardised path coefficients) of the paths in the Structural Equation Model (SEM) used in Fig. 3 (FUNCTION: multifunctionality) and the 16 models using single ecosystem functions (fully-resolved SEMs, Fig. 4) included in the study. Empty cells indicate the weakest insignificant path in the model that was removed in order to calculate SEM model parameters. Env: Environmental variable (PC axis), EW: Earthworm, a: animal, m: microbial, p: plant.

| | | Multifunctionality (see Fig. 3) | | BAS | | $C_{mic}$ | | AMF biomass | | Fungal biomass | | Bacterial biomass | | Grass biomass | | Herb biomass | |
|---|---|---|---|---|---|---|---|---|---|---|---|---|---|---|---|---|---|
| Response | Predictor | p | Path coeff. | p | Path coeff. | p | Path coeff. | p | Path coeff. | p | Path coeff. | p | Path coeff. | p | Path coeff. | p | Path coeff. |
| FUNCTION | EW biomass | **<0.001** | **-0.27** | **<0.001** | **-0.26** | **<0.001** | **-0.29** | 0.95 | 0.01 | 0.06 | -0.35 | 0.16 | -0.25 | - | - | 0.36 | -0.14 |
| FUNCTION | m. multidiversity | 0.13 | 0.10 | 0.26 | 0.08 | **0.04** | **0.14** | 0.24 | 0.16 | 0.91 | -0.02 | - | - | 0.48 | 0.09 | - | - |
| FUNCTION | a. multidiversity | - | - | - | - | - | - | - | - | 0.49 | -0.13 | 0.61 | -0.09 | 0.76 | -0.04 | 0.25 | 0.19 |
| FUNCTION | p. multidiversity | 0.06 | 0.23 | 0.47 | 0.09 | 0.24 | 0.15 | 0.77 | -0.05 | 0.88 | 0.03 | 0.73 | 0.09 | 0.25 | 0.26 | 0.28 | -0.24 |
| FUNCTION | Env1 | 0.42 | -0.15 | 0.82 | -0.05 | 0.14 | -0.30 | 0.19 | 0.28 | - | - | 0.54 | -0.21 | 0.14 | 0.37 | **<0.01** | **0.70** |
| FUNCTION | Env2 | **<0.001** | **0.47** | **<0.001** | **0.58** | **<0.001** | **0.57** | 0.39 | 0.13 | 0.64 | 0.07 | 0.42 | 0.13 | 0.78 | -0.04 | 0.21 | 0.16 |
| m. multidiversity | EW biomass | 0.85 | -0.02 | 0.66 | -0.06 | 0.66 | -0.06 | 0.66 | -0.06 | 0.66 | -0.06 | 0.66 | -0.06 | 0.66 | -0.06 | 0.66 | -0.06 |
| m. multidiversity | Env1 | 0.99 | 0.00 | 0.84 | 0.06 | 0.84 | 0.06 | 0.84 | 0.06 | 0.84 | 0.06 | 0.84 | 0.06 | 0.84 | 0.06 | 0.84 | 0.06 |
| m. multidiversity | Env2 | 0.34 | 0.15 | 0.34 | 0.15 | 0.34 | 0.15 | 0.34 | 0.15 | 0.34 | 0.15 | 0.34 | 0.15 | 0.34 | 0.15 | 0.34 | 0.15 |
| a. multidiversity | EW biomass | **<0.001** | **-0.64** | **<0.001** | **-0.65** | **<0.001** | **-0.65** | 0.34 | 0.15 | **<0.001** | **-0.65** | **<0.001** | **-0.65** | **<0.001** | **-0.65** | **<0.001** | **-0.65** |
| a. multidiversity | Env1 | 0.06 | -0.18 | 0.17 | -0.13 | 0.17 | -0.13 | 0.17 | -0.13 | 0.17 | -0.13 | 0.17 | -0.13 | 0.17 | -0.13 | 0.17 | -0.13 |
| a. multidiversity | Env2 | 0.05 | 0.19 | 0.08 | 0.18 | 0.08 | 0.18 | 0.08 | 0.18 | 0.08 | 0.18 | 0.08 | 0.18 | **0.08** | **0.18** | 0.08 | 0.18 |
| p. multidiversity | EW biomass | **0.02** | **-0.18** | **0.01** | **-0.18** | **0.01** | **-0.18** | **0.01** | **-0.18** | **0.01** | **-0.18** | **0.01** | **-0.18** | **0.01** | **-0.18** | **0.01** | **-0.18** |
| p. multidiversity | Env1 | **<0.001** | **0.66** | **<0.001** | **0.67** | **<0.001** | **0.67** | **<0.001** | **0.67** | **<0.001** | **0.67** | **<0.001** | **0.67** | **<0.001** | **0.67** | **<0.001** | **0.67** |
| p. multidiversity | Env2 | 0.78 | -0.02 | 0.74 | -0.03 | 0.74 | -0.03 | 0.74 | -0.03 | 0.74 | -0.03 | 0.74 | -0.03 | 0.74 | -0.03 | 0.74 | -0.03 |
| Env1 | EW biomass | 0.67 | 0.02 | 0.77 | 0.01 | 0.77 | 0.01 | 0.77 | 0.01 | 0.77 | 0.01 | 0.77 | 0.01 | 0.77 | 0.01 | 0.77 | 0.01 |
| Env2 | EW biomass | **<0.01** | **-0.36** | **<0.001** | **-0.37** | **<0.001** | **-0.37** | **<0.001** | **-0.37** | **<0.001** | **-0.37** | **<0.001** | **-0.37** | **<0.001** | **-0.37** | **<0.001** | **-0.37** |
| m. multidiversity | a. multidiversity | 0.11 | 0.17 | 0.07 | 0.20 | 0.07 | 0.20 | 0.07 | 0.20 | 0.07 | 0.20 | 0.07 | 0.20 | 0.07 | 0.20 | 0.07 | 0.20 |
| m. multidiversity | p. multidiversity | 0.50 | 0.00 | 0.49 | <0.01 | 0.49 | 0.00 | 0.49 | 0.00 | 0.49 | 0.00 | 0.49 | 0.00 | 0.49 | 0.00 | 0.49 | 0.00 |
| a. multidiversity | p. multidiversity | **0.03** | **0.26** | **0.02** | **0.28** | **0.02** | **0.28** | **0.02** | **0.28** | **0.02** | **0.28** | **0.02** | **0.28** | **0.02** | **0.28** | **0.02** | **0.28** |

Table S5 continued.

| Legume biomass | | Woody biomass | | Plant biomass | | Macrofauna biomass | | Mesofauna biomass | | Microfauna biomass | | Soil aggregate stability | | Soil C | | Soil N | |
|---|---|---|---|---|---|---|---|---|---|---|---|---|---|---|---|---|---|
| p | Path coeff. | p | Path coeff. | p | Path coeff. | p | Path coeff. | p | Path coeff. | p | Path coeff. | p | Path coeff. | p | Path coeff. | p | Path coeff. |
| - | - | 0.76 | -0.05 | 0.07 | -0.34 | 0.38 | 0.15 | 0.41 | -0.15 | 0.06 | -0.22 | 0.73 | -0.06 | **<0.001** | **-0.24** | **<0.001** | **-0.26** |
| 0.26 | -0.14 | 0.58 | 0.06 | - | - | 0.18 | -0.18 | - | - | 0.45 | 0.08 | - | - | 0.22 | 0.08 | **0.04** | **0.13** |
| 0.25 | 0.15 | 0.19 | -0.20 | 0.50 | -0.12 | 0.49 | 0.12 | **0.03** | **0.41** | - | - | 0.36 | -0.17 | - | - | - | - |
| 0.26 | 0.26 | **<0.001** | **0.38** | 0.89 | 0.04 | - | - | 0.83 | 0.05 | 0.21 | 0.25 | 0.25 | 0.30 | 0.53 | 0.08 | 0.19 | 0.16 |
| 0.08 | 0.57 | - | - | 0.71 | -0.13 | 0.07 | -0.35 | 0.64 | 0.12 | 0.61 | 0.14 | 0.74 | -0.12 | 0.12 | -0.31 | 0.38 | -0.16 |
| 0.89 | 0.02 | **<0.001** | **0.54** | 0.66 | 0.07 | **0.03** | **0.33** | 0.06 | -0.27 | **<0.001** | **0.57** | 0.12 | -0.25 | **<0.001** | **0.64** | **<0.001** | **0.60** |
| 0.66 | -0.06 | 0.66 | -0.06 | 0.66 | -0.06 | 0.66 | -0.06 | 0.66 | -0.06 | 0.85 | -0.02 | 0.66 | -0.06 | 0.66 | -0.06 | 0.66 | -0.06 |
| 0.84 | 0.06 | 0.84 | 0.06 | 0.84 | 0.06 | 0.84 | 0.06 | 0.84 | 0.06 | 0.99 | 0.00 | 0.84 | 0.06 | 0.84 | 0.06 | 0.84 | 0.06 |
| 0.34 | 0.15 | 0.34 | 0.15 | 0.34 | 0.15 | 0.34 | 0.15 | 0.34 | 0.15 | 0.34 | 0.15 | 0.34 | 0.15 | 0.34 | 0.15 | 0.34 | 0.15 |
| **<0.001** | **-0.65** | **<0.001** | **-0.65** | **<0.001** | **-0.65** | **<0.001** | **-0.65** | **<0.001** | **-0.65** | **<0.001** | **-0.64** | **<0.001** | **-0.65** | **<0.001** | **-0.65** | **<0.001** | **-0.65** |
| 0.17 | -0.13 | 0.17 | -0.13 | 0.17 | -0.13 | 0.17 | -0.13 | 0.17 | -0.13 | 0.06 | -0.18 | 0.17 | -0.13 | 0.17 | -0.13 | 0.17 | -0.13 |
| 0.08 | 0.18 | 0.08 | 0.18 | 0.08 | 0.18 | 0.08 | 0.18 | 0.08 | 0.18 | 0.05 | 0.19 | 0.08 | 0.18 | 0.08 | 0.18 | 0.08 | 0.18 |
| **0.01** | **-0.18** | **0.01** | **-0.18** | **0.01** | **-0.18** | **0.01** | **-0.18** | **0.01** | **-0.18** | **0.02** | **-0.18** | **0.01** | **-0.18** | **0.01** | **-0.18** | **0.01** | **-0.18** |
| **<0.001** | **0.67** | **<0.001** | **0.67** | **<0.001** | **0.67** | **<0.001** | **0.67** | **<0.001** | **0.67** | **<0.001** | **0.66** | **<0.001** | **0.67** | **<0.001** | **0.67** | **<0.001** | **0.67** |
| 0.74 | -0.03 | 0.74 | -0.03 | 0.74 | -0.03 | 0.74 | -0.03 | 0.74 | -0.03 | 0.78 | -0.02 | 0.74 | -0.03 | 0.74 | -0.03 | 0.74 | -0.03 |
| 0.77 | 0.01 | 0.77 | 0.01 | 0.77 | 0.01 | 0.77 | 0.01 | 0.77 | 0.01 | 0.67 | 0.02 | 0.77 | 0.01 | 0.77 | 0.01 | 0.77 | 0.01 |
| **<0.001** | **-0.37** | **<0.001** | **-0.37** | **<0.001** | **-0.37** | **<0.001** | **-0.37** | **<0.001** | **-0.37** | **<0.01** | **-0.36** | **<0.001** | **-0.37** | **<0.001** | **-0.37** | **<0.001** | **-0.37** |
| 0.07 | 0.20 | 0.07 | 0.20 | 0.07 | 0.20 | 0.07 | 0.20 | 0.07 | 0.20 | 0.11 | 0.17 | 0.07 | 0.20 | 0.07 | 0.20 | 0.07 | 0.20 |
| 0.49 | 0.00 | 0.49 | 0.00 | 0.49 | 0.00 | 0.49 | 0.00 | 0.49 | 0.00 | 0.50 | 0.00 | 0.49 | 0.00 | 0.49 | 0.00 | 0.49 | 0.00 |
| **0.02** | **0.28** | **0.02** | **0.28** | **0.02** | **0.28** | **0.02** | **0.28** | **0.02** | **0.28** | **0.03** | **0.26** | **0.02** | **0.28** | **0.02** | **0.28** | **0.02** | **0.28** |
